# A 3D Human Neuron-on-Chip Platform to Monitor Neuronal Injury Responses

**DOI:** 10.1101/2025.08.06.667201

**Authors:** Ruiping Tang, Charles-Francois Latchoumane, Avi Chopra, Md Marzan Sarkar, Chunki Kim, Nathan Gonsalves, Hsueh-Fu Wu, John C. Sentmanat, Alan Liu, Isha Mhatre-Winters, Aditya Mishra, Andrei G. Fedorov, Jay M. Patel, Nadja Zeltner, Steven L. Stice, Jason R. Richardson, Lohitash Karumbaiah

## Abstract

Traumatic brain injury (TBI) is a major cause of neurological dysfunction and long-term neurodegeneration, yet the intrinsic neuronal contributions to TBI pathophysiology remain incompletely defined. Here, we present a novel *Neuron-on-Chip* microfluidic platform that can be used to mechanically injure mature human prefrontal cortex neurons (hPFCs) embedded in three-dimensional (3D) hydrogels, enabling the study of injury responses in pure neuronal cultures. We assessed real-time calcium dynamics across 13 metrics of single-cell and network activity, revealing a biphasic injury response: an early phase (0.5–24 hr) characterized by excitotoxicity, hyper-synchronized bursting, and network collapse; and a late phase (8 d) marked by sustained depolarization and structural remodeling. Secretome profiling uncovered progressive elevations in extracellular pT181 and total Tau from days 1 to 5 post-injury. Cytokine analyses identified early (24 hr) elevations in IP-10, IL-10, IFNα2, and NCAM, and late increases (8 d) in CXCL9 and MPO, linking neuronal activity changes to stage-specific inflammatory signaling. Immunocytochemistry and immunoblotting confirmed temporally ordered upregulation of calpain-1 and active caspase-3 (days 1–3), phosphorylated Tau (AT8+, days 5–8), and neurofibrillary tangle-like Tau aggregates (NFT+, day 8). These findings establish our platform as a scalable microphysiological model for probing the dynamic cellular and molecular sequelae of neuronal response to injury, offering insights into neurodegeneration and opportunities for therapeutic discovery.

## 1. Introduction

Traumatic brain injury (TBI) is a leading cause of long-term sensorimotor and cognitive impairment, largely driven by disruption of neuronal network function^[1]^. The pathophysiology of TBI is multifaceted, encompassing mechanical damage, oxidative stress, neuroinflammation, and cell death. In addition to these acute effects, TBI is increasingly recognized as a trigger for accelerated aging and cellular senescence ^[2–4]^, and as a significant risk factor for late-onset neurodegenerative diseases, including Alzheimer’s disease (AD)^[5]^. A shared hallmark between TBI and AD is tauopathy, characterized by the pathological misfolding and aggregation of Tau protein. These aggregates can propagate in a prion-like manner across neuronal circuits, amplifying neurodegenerative cascades^[6, 7]^. Notably, Tau accumulation is strongly correlated with cognitive decline, even in the absence of amyloid-β (Aβ) pathology^[8]^. Clinical studies have consistently shown elevated levels of total Tau (tTau) and phosphorylated Tau (pT181) in cerebrospinal fluid (CSF) and extracellular fluid following TBI, with persistent accumulation of Tau aggregates in the brain linked to poor outcomes^[9–13]^. While post-TBI neuroinflammation has been implicated in facilitating Tau aggregation and neuronal internalization, these mechanisms are typically attributed to the actions of astrocytes, microglia, and endothelial cells ^[14, 15]^. The intrinsic capacity of neurons to independently initiate or sustain these processes, remains unclear.

Electrophysiological techniques such as whole-cell patch clamp have been widely employed to assess membrane properties and injury-induced changes at the single-neuron level ^[16, 17]^. However, the low-throughput nature of this technique limits its utility for evaluating large-scale network behavior. In contrast, calcium imaging and multi-electrode array (MEA) technologies facilitate the acquisition of simultaneous multi-neuron recordings and provide spatiotemporal insight into network dynamics. While MEAs lack single-unit resolution and are less suited for 3D cultures, calcium imaging, especially when combined with genetically encoded sensors and optogenetic actuators, has emerged as a powerful alternative for resolving both single-cell and network-level activity with high fidelity^[18]^. These recordings, when integrated with analytical frameworks such as spike estimation^[19]^, waveform analysis^[20]^, and graph theory-based connectivity analysis^[21]^, provide a robust toolkit to probe the functional consequences of neuronal injury.

*In vitro* TBI models serve as an ethical and mechanistically accessible new approach methodology (NAM) platform for studying post-injury neuronal responses, while reducing reliance on animal experiments^[22, 23]^. Two-dimensional (2D) *in vitro* systems have been extensively used for drug screening and therapeutic testing ^[24–27]^, yet they fall short in replicating the biomechanical and cellular complexity of native brain tissue. In contrast, 3D neural organoid culture models offer a more physiologically relevant microenvironment, enhancing cell-cell and cell-matrix interactions, and enabling the modeling of tissue-level shear and compression forces seen *in vivo* ^[28, 29]^. Despite these advantages, neural organoid models have several key limitations. These include the need for long-term cultures to establish organoids, high variability, limited reproducibility between organoids, presence of a necrotic core, cellular heterogeneity, and large size, which prevents the use of live-cell calcium imaging methods for acquiring high-resolution single and multi-unit neuronal activity recordings. A temporally resolved 3D *in vitro* model that overcomes these limitations could reveal mechanistic relationships between neuronal network disruption, inflammation, and neurodegeneration in injured neurons.

To address this need, we developed a novel human Neuron-on-Chip platform that facilitates controlled weight-drop injury of human pluripotent stem cell (hPSC) derived human prefrontal cortex (hPFC) neurons embedded in 3D hydrogels. We monitored the injury progression of weight-drop injured hPFC neurons over a period of eight days, *in vitro*. Neuronal activity was characterized via calcium imaging using 13 quantitative descriptors of single-cell and network function. These functional data were integrated with longitudinal secretome analysis, immunocytochemistry, and western blotting to assess inflammatory signaling and intracellular/extracellular Tau dynamics. The Neuron-on-Chip model is a scalable NAM platform that can be used to determine injury progression, dissect neuronal contributions to TBI pathology, and identify early biomarkers of neurodegeneration.

## 2. Results

### 2.1 Weight-Drop Injury Induces Force-Dependent Neuronal Cell Death in a 3D Human Neuron-on-Chip Model

We engineered a custom 3D human *Neuron-on-Chip* platform for long-term neuronal culture, injury induction, and injury response monitoring of hydrogel-embedded hPFC neurons. A 3D-printed negative mold of the device was used to cast polydimethylsiloxane (PDMS) devices that were bonded to a 35 mm glass-bottom dish (Figure 1A–B, Figure S1A-C). Each PDMS device contained four media reservoirs (R1–R4) that supplied nutrients to two cell-gel chambers (C1 and C2). hPFC neurons were differentiated from hPSCs over a period of 35 days, after which they were encapsulated in Geltrex™ hydrogels and seeded into the cell-gel chambers (Figure 1C). The flexible PDMS shells covering the cell-gel chambers were measured to be 1.87±0.16 mm thick. The underlying cell-gel constructs were ∼ 1 mm and had an effective Young’s Modulus of 145.9±16.92 Pa (Figure S2A). Dynamic mechanical analysis (DMA) under load control mode was conducted to obtain the time-dependent biomechanical response of the cell-gel constructs to weight drop impact. Results revealed changes in the elastic behavior of the cell-gel constructs below 10 Hz, as indicated by higher storage modulus compared to loss modulus (Figure S2B). The apparent crossover of the storage and loss modulus at 20 Hz suggests that the material response becomes increasingly viscoelastic under rapid loading (Figure S2B).

**Figure 1.**
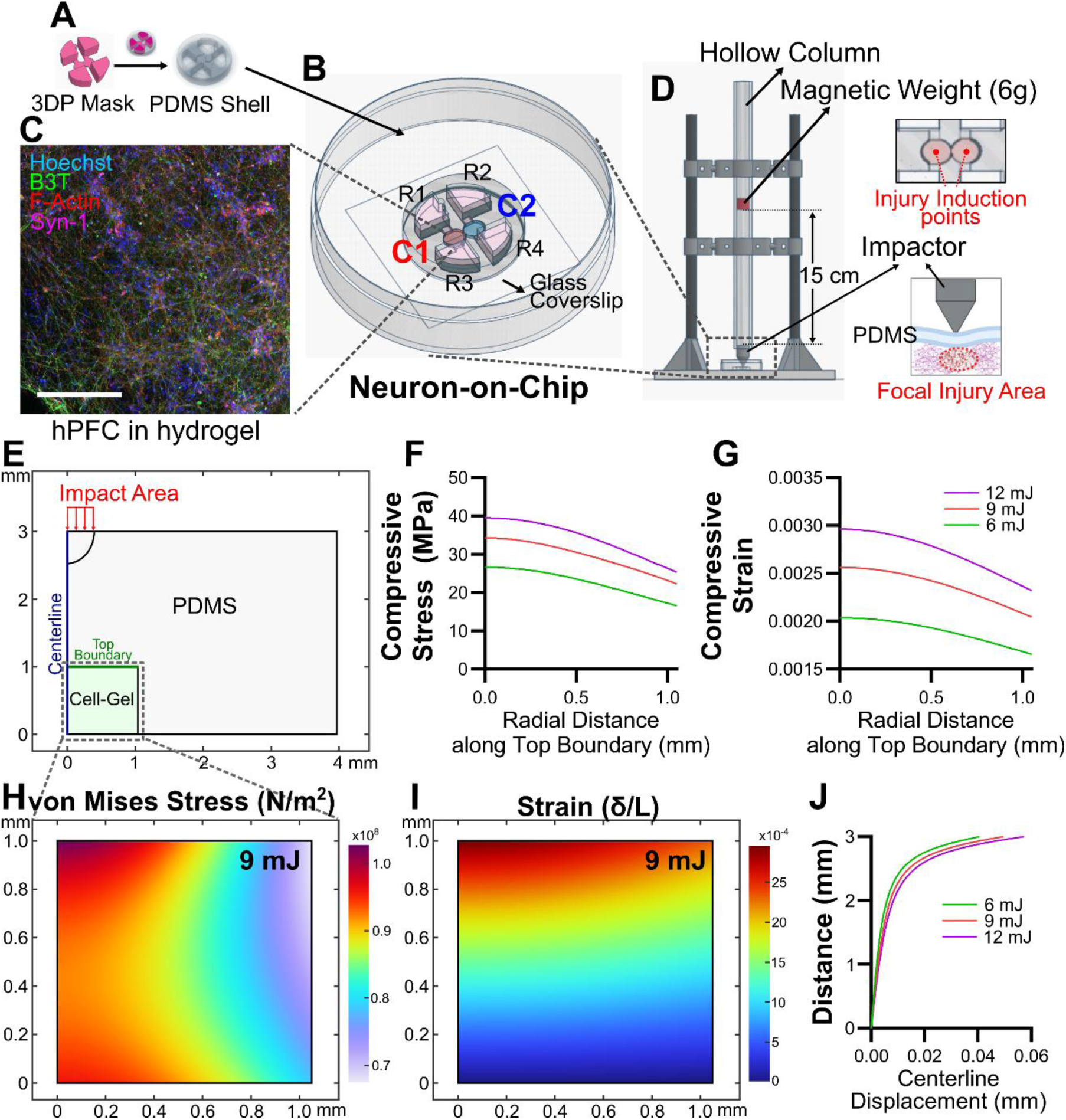
3D Human Neuron-on-Chip device design, fabrication, and characterization. (A) 3D printed negative mold of the device. (B) Schematic of PDMS human Neuron-on-Chip prototype placed on a 35 mm glass-bottom petri dish. Human Neuron-on-Chip device containing hydrogel (Geltrex^TM^) encapsulated hPSC-derived hPFC neurons in reservoirs C1 and C2 that are fed via media reservoirs R1-4. (C) Representative confocal image of immunocytochemically stained mature neurons (Hoechst, blue; β-III-Tubulin, green; F-Actin, red; and Syn-1, magenta). Scale bar – 200 µm. (D) Weight-drop impactor design and setup. (E) 2D-axisymmetric geometry of the PDMS/cell-gel system for FEM simulation. (F-G) Compressive stress (F) and strain (G) vs radial distance along the top boundary (Green line in E) illustrating a graded response across impact conditions. (H-I) Heatmaps of the von Mises Stress (H) and surface strain (I) of the cell-gel construct under 9 mJ injury condition. (J) Displacement of centerline (Blue line in E) indicates the graded response across impact conditions.

After seeding cell-gel constructs into the devices, media exchanges were conducted every 24 hr. After allowing hPFCs maturation for 14 days in the device, immunocytochemical staining and imaging confirmed neuronal network maturation, as indicated by co-expression of the neuronal maturation marker βIII-tubulin and the synaptic protein Synaptophysin-1 (Syn-1), which colocalized with the cytoskeletal marker F-actin (Figure 1C). EdU incorporation assay results indicated minimal proliferation (Figure S3A), supporting neuronal differentiation. Immunocytochemical staining using the pan-neuronal marker HuC/D, astrocyte marker GFAP, and Oligodendrocyte marker Olig2, revealed robust HuC/D+ neuronal enrichment (Figure S3B–C), and minimal glial contamination as determined by GFAP+ and Olig2+ staining, confirming a highly pure neuronal population (Figure S3B–C) 50 days from the start of hPSC differentiation.

We adopted a mechanical weight-drop paradigm to induce focal neuronal injuries using a custom-designed impactor (Figure 1D). A 6 g weight was dropped from 10, 15, or 20 cm heights, delivering ∼6, ∼9, and ∼12 mJ of impact energy, respectively, to a ∼0.5 mm-diameter tip positioned above the neuron-laden hydrogel. The impact transiently deformed the flexible PDMS shell, resulting in localized injury to neurons embedded in the underlying cell-gel constructs. Cell viability was assessed using Hoechst/Calcein AM/Propidium Iodide (PI) staining. Results showed a graded reduction in neuronal survival and neuronal area with increasing impact force (Figure S4A–B). Confocal Z-stack imaging followed by neuronal morphology analyses of Calcein AM stained neurons revealed a significant reduction in soma size, neurite length, and neuronal branch points after injury when compared to no injury controls (Figure S4C-E, Movie S1-2).

To estimate the stress and strain experienced by the neurons upon impact, we analyzed the deformation of the PDMS/cell-gel system using finite element modeling (FEM) following a previously published approach^[30]^, and with geometry and material parameters derived from the Neuron-o-chip device and cell-gel characterization experiments (Figure 1E, Figure S5A). The model predicted that impact-induced deformation was spatially localized to the region beneath the impactor and reached a maximum at approximately 34 µs after contact. Across all the three impact energies, the highest compressive stress and strain occurred along the hydrogel centerline directly beneath the impact site and decayed with radial distance (Figure 1F&G), indicating that the weight-drop delivers a focal mechanical insult rather than a uniformly distributed load. Increasing impact energy from 6 mJ to 12 mJ progressively increased peak cell-gel stress/strain (9 mJ, Figure 1H&I; 6&12 mJ, Figure S5B-E) and centerline displacement (Figure 1J), consistent with the graded reduction in viability and neuronal morphology observed previously.

Based on the graded response observed across injury conditions, we selected the 9 mJ (15 cm drop) as the main injury condition for all subsequent experiments as it produced a clear injury response while preserving sufficient neuronal viability and activity for downstream analyses. Longitudinal assessment of cell viability via Hoechst/PI staining determined that the 6 mJ injury condition produced comparatively milder, and often, insignificant changes in cell viability when compared to uninjured control. The 9 mJ injury condition revealed a modest elevation in PI+ dead cells (<5%) at 0.5 hr post-injury compared to uninjured controls (Figure S4F). A marked increase in cell death (∼10%) was observed between 24 hr and 8 d post-injury (Figure S4F), suggesting a progressive, delayed degenerative response. The 12 mJ injury condition induced a more severe reduction in cell viability but elicited similar neuronal morphology and functional responses compared to the 9mJ condition.

### 2.2 Weight-Drop Injury Elicits Persistent Neuronal Silencing and Biphasic Temporal Network Activity Dynamics

To characterize the effects of mechanical injury on neuronal function, we performed calcium imaging using Fluo-4 AM at five independent timepoints post-injury (0.5 hr, 24 hr, 72 hr, 5 d, and 8 d) across three distinct regions of interest (ROIs). Devices were analyzed for endpoint specific neuronal viability, immunocytochemistry (ICC), western blotting, and cytokine analysis as described in Figure 2A. Due to the need for these endpoint-specific assays, devices allocated for each time point represented an independent terminal sampling point making repeated measurement of the same devices and neuronal populations over time, unfeasible. Calcium imaging confirmed spontaneous spiking activity in hPFC neurons (Figure 2B, Movie S3-6). Neuronal calcium traces were quantified using 13 parameters that represent several single-cell dynamics and network functions, and that were chosen to construct a hierarchical analytical framework to resolve multiple levels of neuronal behavior, ranging from single-cell calcium kinetics, population-level organization, to higher-order network topology and function. These included: Calcium kinetics: mean Rise Time (mRT), Fall Time (mFT), and Bandwidth (mBW), indicative of intracellular calcium handling and burst duration; Signal magnitude: mean Amplitude (Amp), reflecting calcium influx^[31, 32]^; Spike metrics: number of active neurons (AN; ≥5 spikes/min) and mean Firing Rate (mFR), derived from spike estimation algorithms^[20]^. Graph theory was employed to characterize neuronal network structures and identify the presence and connection strength of neuronal subcommunities^[33]^. Network properties: phase-locking value matrices^[34]^ were used to compute weighted Node Degree (wND) and weighted Shortest Path Length (wPL), reflecting cell-to-cell functional connectivity; number of clusters (NoC) and weighted Modularity (wM), representing network community structure; Global Synchronization Index (GSI), indicating the level of synchronized activity; weighted Global Efficiency (wGE), indicating network-wide efficiency; and weighted Clustering Coefficient (wCC), capturing local connectivity patterns^[35, 36]^.

**Figure 2.**
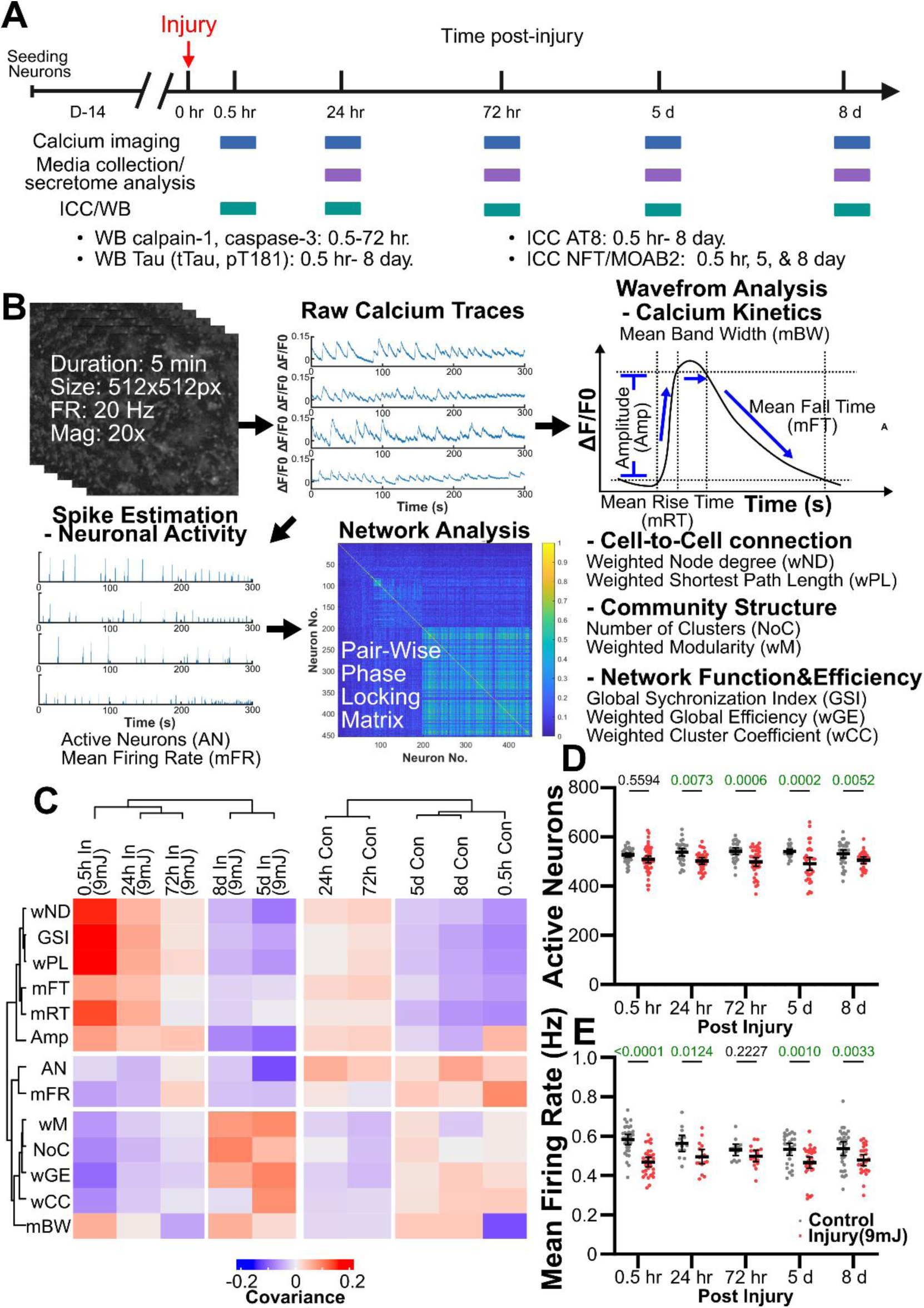
Calcium imaging analysis indicates temporal changes in hPFC neuronal activity after injury. (A) Experiment timeline to assess the temporal effects of weight-drop injury on neuronal activity and network dynamics. Sold blue arrowhead represents the timepoints when the experiments were performed. (B) Schematic of calcium imaging analysis workflow. (C) Heatmap of the results from bi-clustering analysis of 10 sample groups across 13 neuronal function parameters, with distinct neuronal function patterns corresponding to early (0.5-72 hr post injury (In)) and late injury responses (5-8 d post injury (In)). (D, E) Number of active neurons (D) and Mean firing rate (E) of hPFC neurons post-injury. Error bars indicate mean ± 95% confidence interval. Two-way ANOVA with Bonferroni post hoc. P-values of pair-wise comparisons between injury vs control groups at each tested timepoint are reported in the figure and statistically significant differences (p<0.05) are denoted in green text. 3 ROIs per device, 3-5 devices per time point from 4 independent experiments.

Hierarchical bi-clustering of covariance matrices across time and treatment revealed distinct injury-induced patterns (Figure 2C). Compared to uninjured controls, injured neurons showed consistent negative covariance with AN and mFR, suggesting diminished overall activity. AN was quantified to be significantly reduced at all post-injury time points except 0.5 hr (Figure 2D), and mFR was significantly decreased at all time points except 72 hr (Figure 2E), indicating persistent neuronal silencing over the 8-day post-injury period. However, no statistically significant differences in AN and mFR were detected when 9 mJ response was compared to 6mJ and 12 mJ at 0.5 hr and 8 d post injury (Figure S6A-B). Analysis of covariance patterns identified two temporally distinct phases of network response. Early-stage injury (0.5–72 hr) was characterized by positive covariance of wRT, wFT, Amp, wND, wPL, and GSI, relative to controls, whereas these parameters shifted to negative covariance in the late stage (5–8 d). Conversely, NoC, wM, wCC, wGE, and mBW showed opposite trends, demonstrating negative covariance in early injury stages and positive covariance in late injury stages, relative to controls (Figure 2C). Dimensionality reduction via t-distributed stochastic neighbor embedding (t-SNE) and statistical significance test using permutational multivariate analysis of variance (PERMANOVA) analysis revealed significant divergence in multivariate activity profiles between injured and control groups at 0.5 hr, 24 hr, and 8 d, and significant differences between early (0.5-72 hr) and late (5-8 d) injury responses (Figure S7). These analyses support a biphasic trajectory in post-injury network dynamics, delineating early (0.5–72 hr) and late (5-8 d) functional states.

### 2.3 Early Injury Response Is Marked by Synchronized Burst Activity, Dysregulated Calcium Kinetics, and Excitotoxicity Induced Apoptosis

To dissect the early-stage functional response to injury, we analyzed 11 calcium imaging-derived parameters (excluding AN and mFR, which showed global reductions across all post-injury time points) within the first 72 hr post-injury. A hallmark of early injury was significantly elevated global synchronization index (GSI) at both 0.5 hr and 24 hr post-injury relative to uninjured controls (Figure 3A), indicating increased synchronization of neuronal firing.

**Figure 3.**
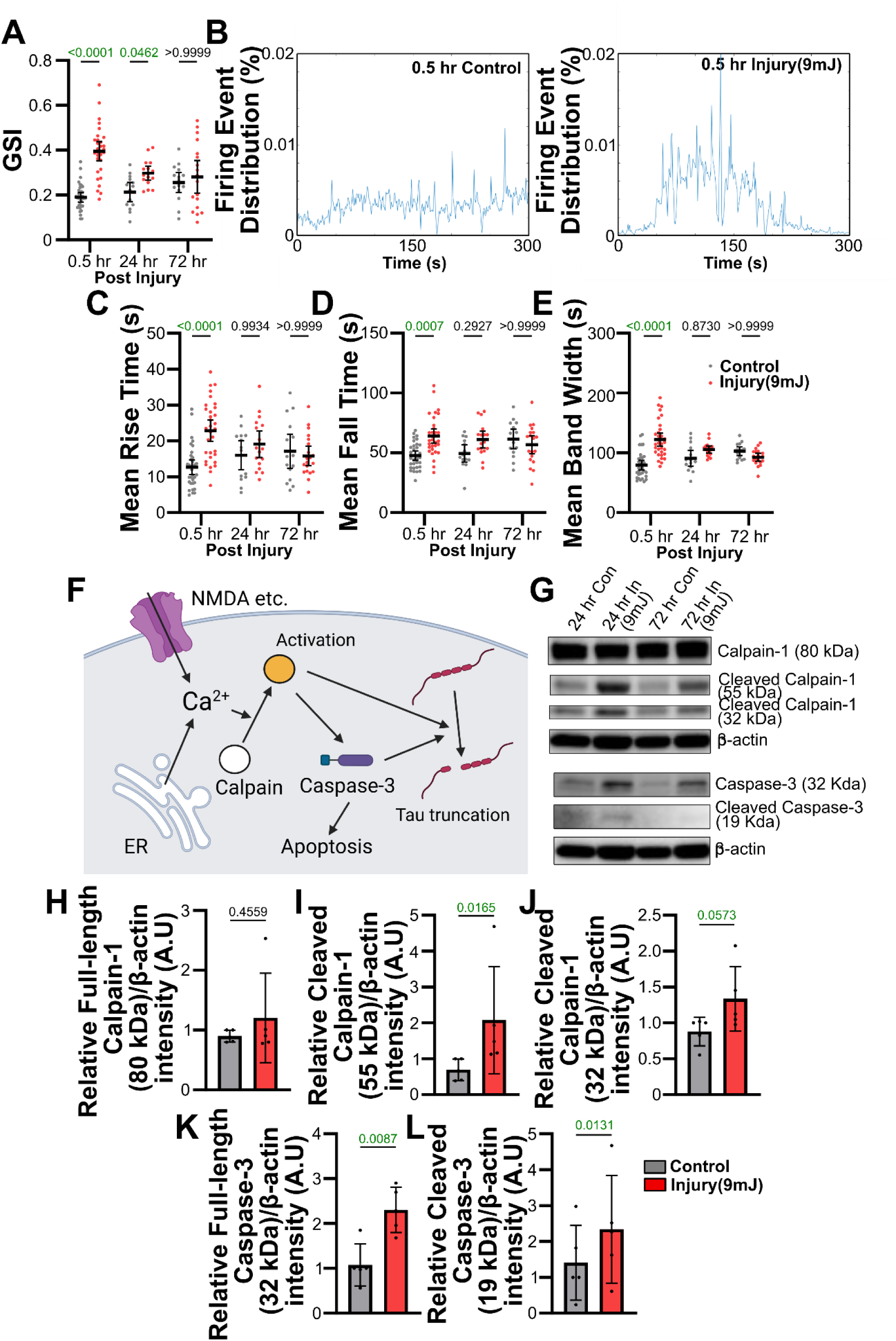
Early injury neuronal response is characterized by highly synchronized burst activity that is driven by excitotoxicity and apoptosis. (A) Global Synchronization Index (GSI) demonstrates the highly synchronized neuronal activity observed acutely post-injury. (B) Firing event distribution from control and injured groups immediately (0.5 hr) post-injury. (C-E) Waveform analyses showing neuronal injury-induced changes in mean rise time (C), mean fall time (D), mean band width (E), acutely, post-injury. Error bars indicate mean ± 95% confidence interval. Two-way ANOVA with Bonferroni post hoc. P-values of pair-wise comparisons between injury vs control groups at each tested timepoint are reported in the figure and statistically significant differences (p<0.05) are denoted in green text. 3 ROIs per device, 3-5 devices per time point from 4 independent experiments. (F) Schematic showing mechanisms of calcium influx induced Calpain activation and downstream effects on caspase-3 and tau truncation. (G) Western blot results of calpain-1 and caspase-3 levels 24-72 hr post injury. (H-L) Quantification of calpain-1 and caspase-3 fragments detected acutely (0.5 hr-72 hr) post injury. Error bars indicate mean ± standard deviation. Welch’s T test, P-values are reported in the figure and statistically significant differences (p<0.05) are denoted in green text. 3-4 devices per sample, 1 sample per time point from 2 independent experiments.

Temporal profiles of calcium traces showed abnormally compact and synchronized burst activity post-injury (Figure 3B), consistent with early hyperexcitability. This was accompanied by significant increases in mean Rise Time (mRT), Fall Time (mFT), and Bandwidth (mBW) (Figure 3C-E), suggesting prolonged intracellular calcium elevation and impaired calcium clearance. Notably, no significant differences were detected in calcium signal amplitude (Amp) during this period, implying that excitotoxic responses were driven by altered dynamics rather than the difference in the magnitude of calcium influx during neuronal depolarization.

At the network level, early injury induced a marked increase in functional connectivity, as evidenced by elevated weighted Node Degree (wND) and Path Length (wPL) (Figure S8A–B; Figure S9). However, this increased connectivity coincided with significant reductions in modularity (wM) and number of functional clusters (NoC) (Figure S8C–E), indicating a collapse of organized network communities. Furthermore, global network efficiency (wGE) was significantly reduced at early time points (Figure S10A), supporting a shift toward a disorganized and energetically inefficient network activity.

To further test if our findings using 9 mJ injury condition can be generalized to other injury conditions, we repeated our calcium imaging experiments at the acute, 0.5 hr post-injury time point for the 6 mJ, and 12 mJ conditions. At 0.5 hr post injury, increasing impact energy produced graded alterations in both calcium kinetics and network organization. Compared to uninjured control, the 9 mJ and 12 mJ conditions showed significantly higher GSI, mRT, mFT and mBW, whereas the 6 mJ condition induced a response that was not significantly different from that of control uninjured neurons (Figure S11A-D). GSI was also significantly higher at 9 mJ and 12 mJ than at 6 mJ, indicating stronger synchronization with increasing injury severity (Figure S11A). In parallel, network community structure became progressively disrupted with increasing impact energy, shown by significant reductions in NoC and wM at 9 mJ and 12 mJ groups compared to the control, with an additional significant decrease at 12 mJ compared to 6 mJ (Figure S11E-F). Similarly, wCC was significantly reduced in 9 mJ and 12 mJ groups compared to the control (Figure S11G), while wGE was significantly rescued in the 12 mJ treatment group compared to both control and the 6 mJ group (Figure S11H). Together, these results indicated that greater impact energies (9 and 12 mJ) induced stronger synchronization, prolonged calcium transients, and pronounced collapse of network communities acutely, with the 6 mJ treatment condition eliciting a response that was not significantly different from controls.

Waveform analysis of neuronal activity patterns suggested that injury-induced excitotoxicity due to excessive calcium influx in excitatory hPFC neurons scaled according to injury severity. Excessive calcium influx is known to activate calpains, a major family of calcium-dependent cysteine proteinases that can contribute to neurodegenerative Tau fragmentation and neuronal apoptosis ^[37, 38]^ (Figure 3F). The patterns of heightened synchronization and prolonged calcium transients observed in our results are consistent with excitotoxic signaling. To assess biochemical correlations of excitotoxic stress, we examined the activation of calpain-1 and caspase-3, two calcium-sensitive proteases implicated in neuronal degeneration. Western blot analysis of cell lysates collected between 0.5 hr and 72 hr post-injury revealed a significant increase in the activated autolytic fragment of calpain-1 (∼55 kDa), with a trend toward elevated 32 kDa fragments (p = 0.0573) (Figure 3G, I–J, Figure S12A-B). Full-length calpain-1 (∼80 kDa) showed only a modest, non-significant increase (Figure 3H).

Consistent with calpain-mediated apoptotic signaling, both full-length (32 kDa) and cleaved (active, 19 kDa) forms of caspase-3 were significantly elevated in injured neurons compared to controls over the same time window (Figure 3G, K–L, Figure S12C-D). These findings support the conclusion that early neuronal hyperactivity leads to calcium overload, triggering calpain-1 autolysis and downstream caspase-3 activation, thereby initiating excitotoxic/apoptotic pathways independent of glial involvement.

### 2.4 Late Injury Response Leads to Sustained Neuronal Depolarization, Network Fragmentation, and Enhanced Community Structure

In contrast to the widespread alterations in neuronal activity patterns observed during the early injury phase, the late-stage injury response (5–8 d post-injury) was characterized by more selective but significant changes in network architecture and calcium dynamics. Notably, injured neurons exhibited sustained depolarization at both 5-d and 8-d post-injury, relative to uninjured controls (Figure 4A), indicating long-term disruption of membrane potential homeostasis.

**Figure 4.**
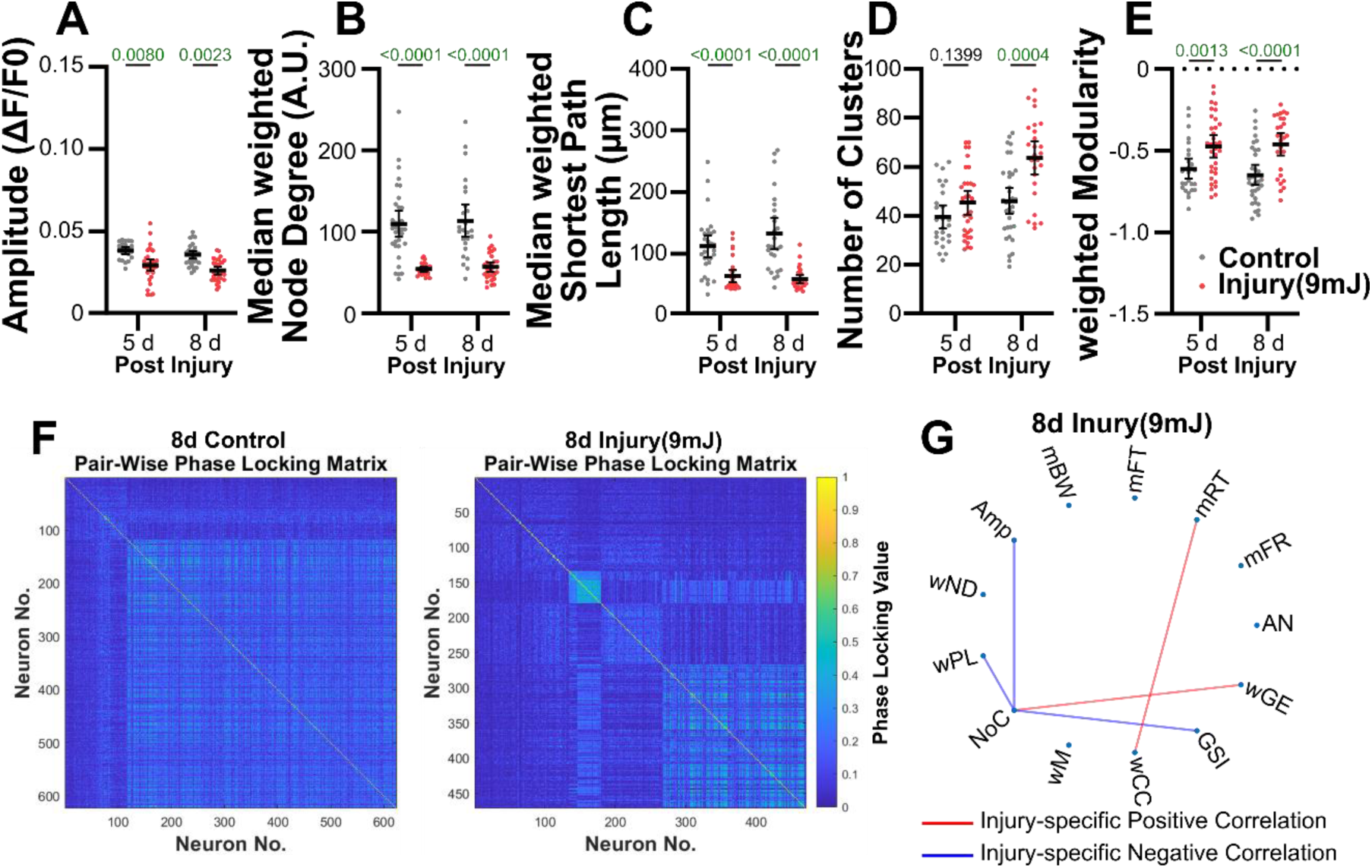
Weight-drop injury induces neuronal depolarization, loss of long-range connectivity, and rearrangement of neuronal community structure. (A) Amp of calcium signal demonstrates neuronal depolarization 5-8 d post injury. (B, C) Median weighted Node Degree and median weighted Shortest Path Length reveal acute increase and long-term loss of functional connections. (D, E) Number of Clusters (D, neuronal subcommunities) identified by an eigen-value-based algorithm, and weighted Modularity (E, presence of neuronal sub-communities) demonstrate acute disruption and eventual rearrangement of neuronal community structure in response to injury. Error bars indicate mean ± 95% confidence interval. Two-way ANOVA with Bonferroni post hoc. P-values of pairwise comparisons between injury vs control groups at each tested timepoint are reported in the figure and statistically significant differences (p<0.05) are denoted in green text. 3 ROIs per device, 3-5 devices per time point from 4 independent experiments. (F) Representative heatmaps of pair-wise phase locking matrix from control and injured groups 8 d post injury, reveal the remodeling of functional network structure in response to the injury. (G) Differential network analysis of injured vs control groups, 8-d post-injury reveals injury specific correlation between NoC, AmP, wPL, and GSI.)

At the network level, injured neurons showed significant reductions in wND and wPL at late time points (Figure 4B&C; Figure S9), indicating weakened connectivity and loss of long-range functional links. Concurrently, an increasing trend in NoC was observed, with significant elevations at 8 d post-injury (Figure 4D). wM also increased significantly at 5 and 8 d (Figure 4E–F), suggesting reorganization of neurons into smaller, more segregated communities. These changes were partially accompanied by a rising trend in wGE (Figure S10A) and a statistically significant increase in wCC at 5 d post-injury (Figure S10B), reflecting enhanced local neighborhood connectivity. Additionally, minor but consistent upward trends in mRT, mFT, and mBW were observed in injured neurons over this period (Figure S13B–D), suggesting that mild dysregulation in calcium kinetics remained even at later time points. When comparing across the injury conditions, we found stress-dependent response and significant reduction of Amp in 12 mJ vs 6 mJ conditions (Figure S14A). However, similar responses was not observed in wM and NoC (Figure S14B-C).

Biochemical markers of cell death and excitotoxic stress diverged from the early-stage profile. Calpain-1 cleavage products remained unchanged compared to controls at late time points (Figure S12A-B), and while full-length caspase-3 levels significantly decreased in injured neurons at 5–8 d, the active cleaved form (19 kDa) was no longer detectable (Figure S12C-D). These findings suggest a resolution or exhaustion of the apoptotic response in the late injury phase.

To further probe structural changes in network topology, we performed differential network analysis^[39]^ restricted to the 8-day post-injury time point—chosen due to the significant divergence between injured and control groups across all 13 functional parameters as confirmed by PERMANOVA. This analysis revealed that NoC functioned as a central network hub that negatively correlated with Amp, wPL, and GSI, and positively correlated with wGE (Figure 4G). These results indicate that injury-induced fragmentation of large-scale connectivity is accompanied by a compensatory increase in modular community structures, and that community reorganization is closely linked to signal synchronization, calcium amplitude, and network efficiency.

### 2.5 Secretome Analysis Reveals Extracellular Tau Accumulation and Temporal Regulation of Inflammaging Associated Proteins Post-Injury

To identify injury-induced changes in secreted neurodegenerative and inflammatory markers, we analyzed the conditioned media from injured and uninjured hPFC neuron cultures over four timepoints: 24 hr, 72 hr, 5 d, and 8 d post-injury. Log₂ fold changes were computed to compare the abundance of secreted proteins between injured and control groups.

Among neurodegeneration-related proteins, we observed a sustained increase in extracellular phosphorylated Tau (pT181) across the first 5 days post-injury in the injured group, indicating persistent release of pathologically modified Tau isoforms (Figure 5A). This was accompanied by elevated levels of total Tau, suggesting broad Tau dysregulation in response to mechanical insult. In contrast, extracellular levels of Aβ40 and Aβ42 progressively declined post-injury, diverging from Tau trends and suggesting distinct regulatory mechanisms governing amyloid and Tau dynamics in neuron-only cultures (Figure 5A). Temporal clustering of 24 inflammatory and neurodegeneration-associated proteins revealed three secretion profiles: 12 proteins exhibited early release (peak at 24 hr), 8 proteins showed mid-phase release (peak at 5 d), and 4 proteins displayed late release (peak at 8 d), indicating a structured temporal cascade in the neuronal secretome post-injury.

**Figure 5.**
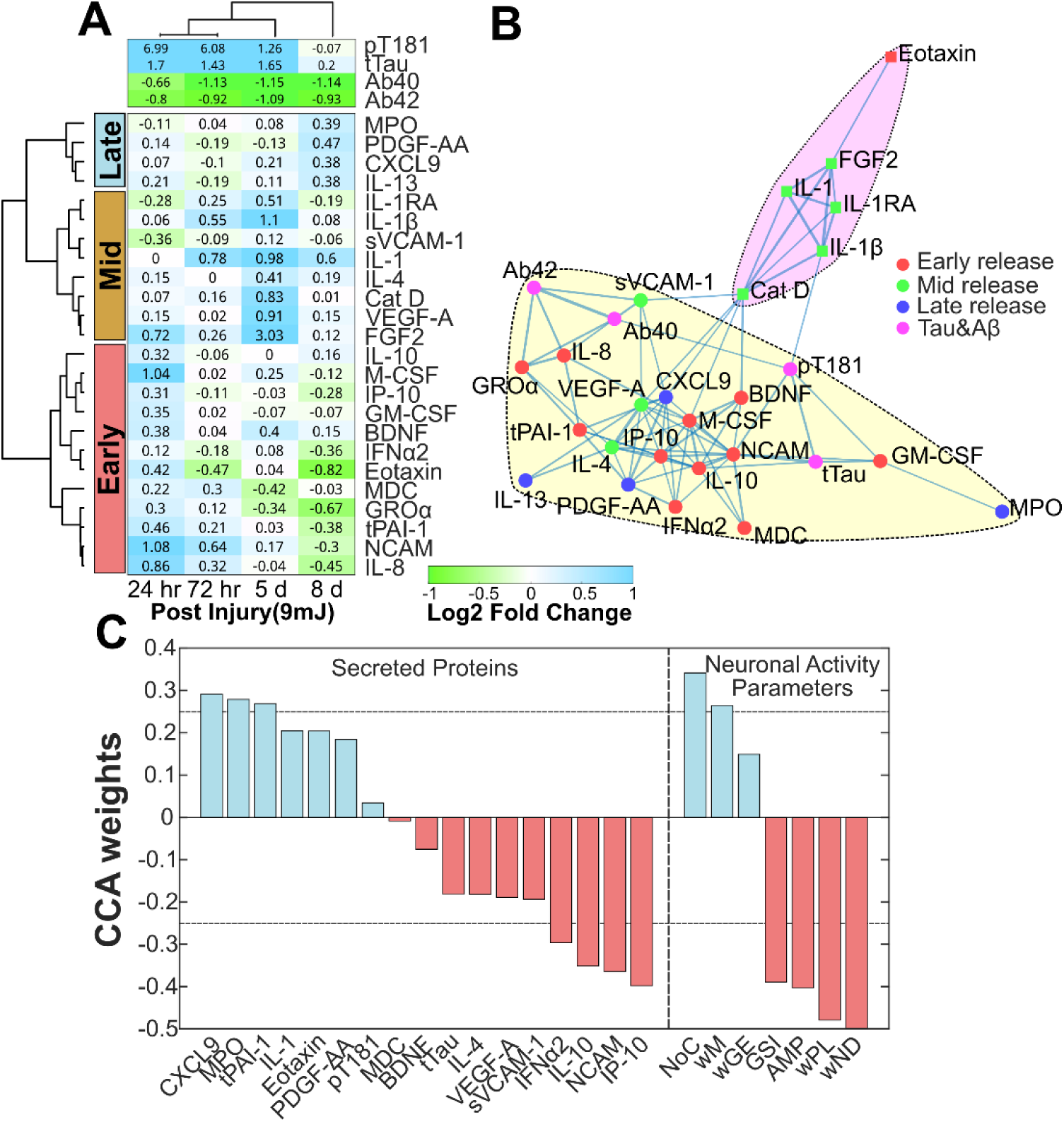
Secretome analysis of inflammatory and neurodegenerative proteins validates extracellular Tau localization, and shows temporal regulation of inflammatory factors, post-injury. (A) Log_2_ fold change of secreted proteins in injured vs control groups across time indicates temporal regulation and classification into three clusters - Early, Mid, and Late-release. 3-5 devices per sample, 1 sample per timepoint from 2-3 independent experiments. (B) Network analysis and spectral clustering analysis reveal 2 functional clusters within 28 secreted factors and their hub proteins. (C) Sparse canonical correlation analysis identified the high association of IP-10, IL-10 and NCAM with early injury state neuronal function; and CXCL9, MPO, tPAI1, with late injury state neuronal function.

Spectral clustering of protein-protein interaction networks identified two major clusters with distinct functional and temporal signatures (Figure 5B). The yellow cluster (early-phase hub) contained IP-10, IL-10, NCAM, and M-CSF—factors that formed central nodes connected to both mid-phase (IL-4, VEGF-A) and late-phase factors (CXCL9, PDGF-AA). The purple cluster included predominantly mid-release proteins, such as FGF2 and IL-1 family cytokines, which also showed the highest fold changes across all time points. Cathepsin D (CatD) was identified as a key regulator bridging interactions between the two clusters.

To explore relationships between secretome dynamics and functional neuronal outcomes, we performed sparse canonical correlation analysis (CCA) ^[40]^ using the 13 calcium imaging-derived activity parameters (Figure 5C). CCA separated these parameters into two functional groups: Group 1 (Red): Early injury-associated metrics, including AMP, GSI, wND, and wPL; and Group 2 (Blue): Mid-to-late-stage parameters, including wGE, NoC, and wM. Secreted proteins strongly correlated with Group 1 included early-phase cytokines IP-10, IL-10, NCAM, and IFNα2 (CCA weight > 0.25), indicating tight association with initial network hyperactivity and connectivity changes. In contrast, Group 2 activity parameters correlated most strongly with late-phase proteins such as CXCL9 and MPO, along with tPAI1, which peaked earlier but exhibited delayed associations with network remodeling. Additional secreted factors-VEGF-A, IL-4, BDNF, MDC, pT181 Tau, tTau, PDGF-AA, Eotaxin, and IL-1 demonstrated significant correlations with neuronal activity but were assigned lower CCA weights (<0.25), suggesting more modest or indirect contributions to functional dynamics.

Together, these findings indicate that Tau release and injury-triggered inflammatory signaling are temporally structured, and that secreted factors—particularly those in the early-phase yellow cluster—are tightly coupled to specific features of network activity and remodeling. This reinforces the functional interdependence between neuronal secretome composition and activity state following mechanical injury.

### 2.6 Weight-Drop Injury Induces Intracellular Tau Aggregation that is Correlated with Neuronal Activity Disruption

Given the persistent elevation of extracellular total Tau and phosphorylated Tau (pT181) identified in the secretome analyses, we next investigated intracellular Tau dynamics in injured hPFC neurons using western blotting and immunocytochemistry (ICC).

Western blot analysis revealed a significant reduction in intracellular tTau (total Tau, Tau46) at 24 hr, 5 d, and 8 d post-injury compared to uninjured controls (Figure 6A&B), suggesting enhanced Tau release or degradation following neuronal injury. In contrast, levels of intracellular phosphorylated Tau (pT181) were significantly elevated across all time points post-injury (Figure 6A&C), consistent with abnormal Tau phosphorylation and aggregation. Other phospho-Tau species, including pTau231 and pS396, were also examined. While pTau231 showed no significant change, pS396 displayed a trend toward upregulation in injured neurons (p = 0.0594; Figure S15), indicating potential isoform-specific involvement in injury-induced Tau pathology.

**Figure 6.**
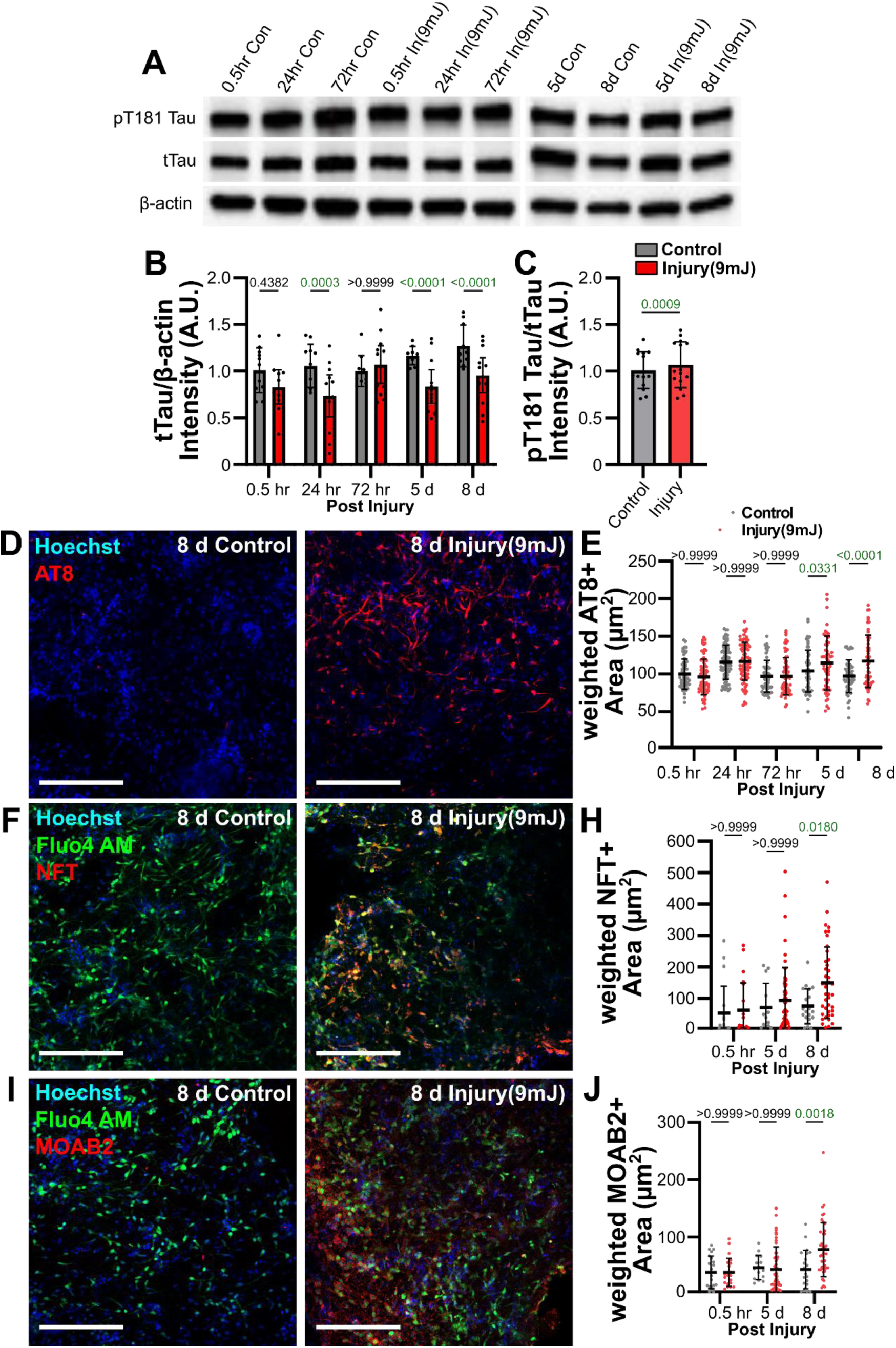
Weight-drop injury induced the intracellular accumulation of pathological Tau and Aβ isoforms in hPFC neurons seeded in 3D Neuron-on-Chip devices. (A) Western blots of pT181 Tau, total Tau (Tau46) and β-actin loading controls from injured and control groups across time points. (B-C) Quantification of results from (A). 3-4 devices per sample, 1 sample per time point from 3 independent experiments. Total Tau measurement was repeated 4 times per sample from each independent experiment. (D) Representative confocal image of immunocytochemically stained AT8 (pathological Tau maker, red). Scale – 200 µm. (E) Quantification of AT8 area demonstrating significant AT8 upregulation 5- and 8-day post injury in injury groups, compared to the control. (F) Representative confocal image of Fluo4-AM immunocytochemically stained NFT (neurofibrillary tangles, red). Scale – 200 µm. (G) Quantification of NFT area demonstrating significant NFT upregulation 8-day post injury in injury group, compared to the control. (H) Representative confocal image of Fluo4-AM and immunocytochemically stained MOAB2. (I) Quantification of MOAB2 area demonstrating significant MOAB2 upregulation 8-day post injury in injury group, compared to the control. Scale – 200 µm. Error bars indicate mean ± standard deviation. 8-10 images per device, 3 devices per group, from 2 independent experiments. Two-way ANOVA with Bonferroni post hoc. P-values of pair-wise comparisons between injury vs control groups at each tested timepoint are reported in the figure and significant p-value (<0.05) is highlighted in green.

To further characterize Tau and Aβ aggregation and maturation, we performed ICC using antibodies against hyperphosphorylated Tau (AT8; paired helical filament-positive), neurofibrillary tangle (NFT)-positive Tau, and MOAB2 positive Aβ (unaggregated, oligomeric, or fibrillar forms of Aβ42 and unaggregated Aβ40). Staining results revealed significant increases in AT8+ Tau in injured neurons at 5-d and 8-d post-injury (Figure 6D&E). Additionally, NFT+ Tau and MOAB2+ Aβ was markedly elevated at 8-d in the injured group compared to controls (Figure 6F-J), confirming the late-stage accumulation of aggregated Tau and Aβ structures. We further compared tau progression across 3 injury conditions (6 mJ, 9 mJ, and 12 mJ) using antibody against AT8. Although all three impact energies tested significantly enhanced AT8+ Tau in injured neurons, no significant differences in AT8+ Tau expression were detected between injury conditions (Figure S16).

These findings establish a clear link between injury-induced changes in neuronal activity and the intracellular accumulation and aggregation of pathogenic Tau species. These results also provide evidence of extracellular Tau release dynamics and demonstrate the progressive development of tauopathy in weight drop injured pure neuronal cultures.

## Discussion

Neuronal injury is characterized by acute neurotoxicity and widespread disruption of activity patterns, often resulting in the rewiring of neuronal circuits. While these findings have been well documented in vivo, and in mixed cell culture models, the direct implications of injury and injury-induced neurodegeneration in genetically unaltered, neuron-only cultures remain poorly understood. Using a 3D hydrogel-based Neuron-on-Chip platform incorporating hPFC neurons, we demonstrate that mechanical injury via weight-drop leads to temporally dynamic changes in both single-unit and network-level activity. These activity changes are accompanied by a tightly regulated and temporally distinct inflammatory secretome, which together activate neurodegenerative tau pathology.

Immediately post-injury, neurons exhibited hyperexcitation driven by glutamate-induced excitotoxicity and the acute loss of inhibitory input-hallmarks of TBI pathology^[41, 42]^. By modeling injury responses using a purified excitatory hPFC neuronal population, we aimed to isolate cell-intrinsic mechanisms of damage response. These hPSC-derived neurons are predominantly cortical projection neurons^[43]^, which are well-suited for modeling neurodegenerative responses^[44–46]^. Using Fluo-4 AM calcium imaging, we quantified intracellular Ca^2+^ transients and derived event/spike estimates and network function metrics from deconvolved traces. Calcium-based metrics provide a robust comparable readout of neuronal activity across experimental groups. Additionally, we interpret firing activity and network structure as inferred activity measures, providing insights beyond the temporal and spatial limitations of MEA or patch clamp approaches.

Our results reproduced classical excitotoxic responses to injury, including calcium dyshomeostasis and synchronized burst firing, that scales with injury severity. These findings are consistent with previous work reporting increased membrane permeability^[47, 48]^, dysregulation of voltage-gated Na⁺ channels^[49, 50]^, and glutamate-induced excitotoxicity^[51]^, and also suggesting more extensive membrane perturbation and stronger excitotoxic signaling resulting from greater mechanical loading. These processes provide a plausible basis for the higher GSI and the prolonged calcium kinetics observed at 9 mJ and 12 mJ conditions, as greater calcium dysregulation would lead to higher neuronal activity and slower calcium trafficking kinetics ^[52]^. The increased cleavage of calpain-1 observed via western blotting studies further confirmed calcium dysregulation. Calpain, a calcium-activated protease, undergoes autolysis in the presence of elevated Ca²⁺ levels^[53]^, thereby facilitating excitotoxic damage by proteolyzing voltage-gated Na⁺ channels^[54, 55]^ and promoting tauopathy.

Functional connectivity analyses revealed that early synchrony post-injury was followed by significant reduction in connectivity strength, depolarization, and reorganization of neuronal clusters. These late-phase changes suggest long-term depression (LTD)-like plasticity, potentially mediated by decreased calcium influx, a known trigger for shifting synaptic strength from long-term potentiation (LTP) to LTD^[56, 57]^. The reduced network amplitude observed aligns with these LTD-associated mechanisms, characterized by lower wPL and increased community fragmentation. These data present a comprehensive view of the temporal evolution of neuronal injury responses across both cellular and network dimensions.

To investigate molecular underpinnings of these functional transitions, we profiled secreted proteins across injury time points. While neuronal production of inflammatory cytokines is known^[58–75]^, our study provides one of the first high-resolution temporal maps of cytokine secretion from pure neuronal populations. We identified 24 secreted proteins, many of which play roles in regulating neuronal function and inflammatory signaling^[69, 70, 76–82]^. Secretome dynamics revealed temporally defined phases of release: early (e.g., PAI-1, BDNF, IL-8, IP-10), mid (e.g., IL-1 family, PDGF-AA), and late-phase (e.g., IL-13, MPO) factors. Notably, many of these proteins are also upregulated within 3 days post-TBI in both human and animal studies^[83–87]^.

We propose that early-stage excitotoxicity and hyperactivity likely drive the release of activity-regulated factors, as suggested by high canonical correlations between IP-10, IL-10, IFNα2, NCAM, and injury-induced activity changes. IP-10 and NCAM, in particular, are upregulated via calcium-dependent gene expression mechanisms^[88–90]^, and the involvement of CDK5 and calpain in tau hyperphosphorylation have been well established. While direct activity-dependence of IL-10 and IFNα2 remains uncertain, their tight association with early injury responses suggests indirect regulation through other neuronal signaling cascades. Network analysis further revealed IP-10, NCAM, IL-10, and M-CSF as central hubs in autocrine regulatory circuits, indicating a mechanistic link between neuronal activity and inflammatory secretome progression.

We found that several injury-induced secreted proteins are components of the senescence-associated secretory phenotype (SASP), which exerts autocrine and paracrine effects in aged tissues^[91–95]^. These include early-phase proteins (e.g., IL-8, GM-CSF), mid-phase (e.g., FGF2, IL-4), and late-phase (e.g., IL-13, VEGF-A). In parallel, we identified non-SASP factors such as MPO and Cathepsin D, which have been directly linked to tau pathology^[15, 58, 96–100]^. Together, these findings suggest that neuronal injury alone can recapitulate aging-like secretory patterns and drive neurodegenerative changes, independent of glial involvement. These molecular profiles coincide temporally with the observed shift from early hyperexcitability to late-phase LTD, supporting existing links between aging, synaptic weakening, and tau aggregation^[101–104]^.

The extracellular release of neurodegenerative Aβ was decreased following injury, consistent with human CSF profiles post-TBI^[105, 106]^. This reduction may reflect enhanced intracellular accumulation of Aβ due to upregulated APP processing via injury-induced neuronal activity^[107–110]^, and is supported by our findings of delayed MOAB2 accumulation. In contrast, extracellular tau (total and pT181) increased significantly, confirming previous TBI findings^[9, 10]^. The increase in extracellular Tau along with simultaneous decline in intracellular total Tau and rise in intracellular pT181 suggests accumulation of hyperphosphorylated tau species that potentially contribute to the formation of tau aggregates and neurofibrillary tangles. Moreover, our results suggest a temporally structured sequence in which functional disruption and neuronal Tau pathology evolve in parallel. The timing of these changes is consistent with bidirectional coupling rather than a single unidirectional pathway, supported by prior reports demonstrating that neuronal activity can modulate Tau secretion^[111–114]^, and that pathological Tau leads to synaptic impairment, network communication destabilization and LTD^[115–119]^. Although, significant differences in Amp were observed at 8-d post injury across the injury conditions tested, no significant differences in network level activity as well as cellular Tau readouts were observed, suggesting that the tested injury conditions could not resolve stress-dependent changes in network level and Tau deposition. Future studies could test the effects of higher impact forces and pharmacological rescue experiments on changes in network structure/function and Tau readouts.

Calpain-1, cathepsin D, and active caspase-3 enzymes that are known to cleave tau at specific sites and generate neurotoxic fragments^[120–122]^, were found to be significantly upregulated after neuronal injury. Tau truncation products such as Tau_151-391_ have been shown to exhibit high aggregation propensity and pathological phosphorylation^[123]^. Calpain-1 and caspase-3 are primarily responsible for the formation of tau fragments ^[120, 124]^. Calpain-1 produces a cleaved form of ∼ 78 kDa ^[124]^ and activated forms at ∼ 55 and 32 kDa ^[120, 125–127]^ by autolysis. We observed that these activated forms of calpain-1 were elevated post-injury followed by increased active caspase-3 early in injury. Indeed, previous studies have shown that calpain-1 knockout mice lack the ability to increase caspase-3 and its active form after injury ^[38]^, suggesting that calpain-1 directly produces active caspase-3 in our model. Additionally, the total Tau antibody Tau46, detects AA404-441 on the C-terminal of Tau protein, which can be cleaved by the three proteinases. The significant decrease of intracellular Tau46+ staining post-injury is consistent with enhanced tau truncation.

A variety of in vitro TBI models have been previously developed to reproduce different aspects of injury biomechanics and cellular response ^[28, 29, 128]^. These include conventional 2D stretch/shear/blast paradigms using dissociated neuronal cultures ^[48, 129]^, organotypic brain slices ^[130, 131]^ that enable controlled cellular compartmentalization, and 3D hydrogels and brain organoid approaches that approximate tissue-like cell-matrix interactions ^[132–151]^. Recent reviews highlight that 2D models are often well-suited for high-throughput screening but can underrepresent 3D biomechanical properties, whereas organoids and 3D cultures improve physiological translatability but can introduce variability, limit optical access, and require prolonged culture time ^[29, 130, 152]^. Existing 3D in vitro TBI models include multicellular organotypic slices ^[130, 131, 141]^, brain organoid ^[141, 143, 146]^, and engineered 3D tissue systems ^[142, 148, 150, 151]^ that capture neuron-glia interactions. However, except for organotypic tissue and neurovascular-unit models ^[145]^, the vast majority lack vascular components. While the Neuron-on-Chip model also lacks glial and vascular components, it demonstrates the innovative use of this platform to investigate neuron only responses to injury. The device design and fabrication schemes significantly reduce the complexity of device manufacturing by eliminating the need for advanced cleanroom and soft photolithography facilities. Our approach facilitates greater throughput, dynamic imaging, media accessibility, and scalability to include other cells and therapeutic factors, which are advantageous features for perturbation and therapeutic screening studies. By subjecting 3D hydrogel embedded hPSC-derived hPFC neurons in Neuron-on-chip devices to weight drop injury, we demonstrate the manifestation of pathophysiologically relevant acute excitotoxicity, progressive loss of functional connectivity, and the emergence of a neurodegenerative and SASP-like secretome, in relevant excitatory neuron populations, and in the absence of glial contributions. To our knowledge, these findings represent the first direct association between injury induced neuronal activity trajectories and inflammatory/neurodegenerative biomarkers in a neuron-only setting. These findings also highlight an intrinsic neuronal mechanism for tauopathy development following neuronal injury and offer a unique platform to dissect cell-autonomous injury responses relevant to TBI and neurodegenerative disease progression.

## 4. Limitations of the Study

While the 3D Neuron-on-Chip model offers a scalable and highly controlled *in vitro* platform to study injury-induced neuronal responses, several limitations are acknowledged. First, the system lacks glial and vascular components, which play critical roles in modulating neuronal excitability and inflammatory signaling. Although the neuron-only experimental design setup enabled the dissection of intrinsic neuronal injury responses, the broader contribution of neuroglial and vascular endothelial cells were not assessed in this study. The use of a cell-permeable calcium indicator dye (Fluo-4 AM) limited our ability to longitudinally track the activity of the same neuronal population over time. As devices were sampled at terminal endpoints, temporal differences should be interpreted as population-level changes between treatment groups rather than longitudinal within-device progression. Calcium imaging is an indirect measure of neuronal activity and does not directly quantify action potentials or synaptic currents. Electrophysiological validation of calcium responses and extension of findings across multiple donor cell lines remain to be determined. While device fabrication and injury applications can be performed in batches, live calcium imaging and downstream imaging analysis are currently the most time-intensive steps because each device must be imaged individually under standardized acquisition conditions. Future work will address these limitations by including neuroglial cells, additional donor cell lines, implementing genetically encoded calcium indicators (e.g., GCaMP) to facilitate time course changes within the same device, and implementing concurrent electrophysiological validation.

## 5. Methods

*Device Fabrication:* Reusable negative device molds were 3D printed using Clear-V4 resin (Formlabs™, Cat# RS-F2-GPCL-04) on a Formlabs 3 printer. PDMS (Sylgard™ 184, Dow Corning) was mixed in a 10:1 base-to-curing agent ratio and then degassed under vacuum for 0.5 hr - 1 hr. Then 7 mL of the mixture was poured into 60 mm Petri dishes containing seven molds that were held in place with mineral oil. Device molds and PDMS mixture were degassed under vacuum again for 0.5 hr - 1 hr. To ensure consistency in PDMS thickness and to reduce batch to batch variation, petri dishes containing PDMS and 3D printed molds were placed on a level surface in the oven for 30 mins before being cured at 80°C for 3 hours to result in polymerized PDMS devices that were ∼3 mm in height. Polymerized PDMS using the above ratio of base polymer and curing agent has been previously characterized to have an elastic modulus of ∼1.3-3 MPa^[153]^. The PDMS devices were removed from 60 mm petri dish using a 13 mm diameter biopsy punch and inspected for any visible bubbles, incomplete curing or defects around the culture chamber. The devices were subsequently immobilized on clean 35 mm glass bottom dishes (Cellvis) using uncured PDMS base polymer. The devices were returned to the oven to bake for another 3 hours at 80°C, then sterilized in 70% ethanol and air-dried before seeding cells. The frequency of device fabrication is determined by the number of molds obtained from each batch of 3D printing. In general, 48-60 molds were used to produce a single batch of PDMS devices with less than 5% of devices failing quality check. The major causes of device exclusion are visible PDMS defects including bubbles or uneven cropping using the 13 mm diameter biopsy.

*Cell Culture of hPSC-Derived Prefrontal Cortex Neurons:* Human pluripotent stem cells (hPSCs) derived prefrontal cortex neurons (hPFCs) were generated following methods described previously ^[43]^ using Mel1 cell line (NIHhESC-11-0139, passage number 40-50). PFCs were used for experiments 35 days after differentiation. Detailed procedure is included in the supplementary method.

*Hydrogel encapsulation of hPFC neurons:* hPSC-derived hPFC neurons were suspended in Geltrex^TM^ at a concentration of 20,000 cells/μL. 10 μL of mixture was quickly dispensed into each cell chamber (B1, B2) of the device. The device was then incubated at 37°C/5% CO_2_ for 30 minutes to facilitate polymerization before 200 μL of PFC media was added into the device. Media change was performed daily thereafter. Geltrex^TM^ is a basement membrane extracellular matrix extract that is primarily composed of laminin, collagen IV, entactin/nidogen, and heparan sulfate proteoglycans. Gelation occurred through the intrinsic thermoresponsive self-assembly of Geltrex^TM^ upon incubation at 37 °C. We found that variability in hydrogel formation stemmed from lot-to-lot differences in Geltrex^TM^. To minimize this source of variation, the same lot of Geltrex^TM^ was used for each batch of experiments as much as possible. Because hPSC-derived PFC neurons include populations of undifferentiated proliferative cells during the initial phase after seeding, we consistently utilized EdU incorporation assay, standardized cell passaging, and seeding and assessment timelines to maintain a uniform density of neuronal cells in all devices.

*Weight-Drop Injury Induction:* hPFC neurons in the device were allowed to mature for 2 weeks before being subjected to weight-drop injury. A weight drop apparatus was designed and built based on Feeney’s weight drop injury model ^[154, 155]^. Both chambers within each Neuron-on-Chip device received a focal injury with a 6-gram weight dropped from a height of 15 cm (impact velocity ≈ 1.72 m/s, acceleration ≈ 1 g, force ≈ 9 mJ, force/area on PDMS shell ≈ 11.4 mJ/mm^2^).

*Mechanical characterization of the hPFC-hydrogel mix:* The stiffness characterization of cell-gel construct was done by indentation method with Pavone nanoindenter (Optics11 Life). hPSC-derived hPFC neurons were suspended in Geltrex^TM^ at a concentration of 20,000 cells/μL. The hydrogel encapsulating cells was plated in a 24-well plate prior to the measurements. Indentations were done with a spherical probe of 26.5 µm tip radius and stiffness of 0.015 N/m, which was pre-calibrated by the manufacturer. The effective Young’s modulus was calculated from Hertzian contact model and fit up to 10 nN of the force curve. Dynamic mechanical analysis (DMA) was performed by setting have an initial 40 second relaxation at 0.1 uN, followed by a 10 nN oscillation at 1, 2, 5, 10, and 20 Hz with a 2 second relaxation between each frequency.

*Finite element modeling (FEM):* The simulation of the deformation of the PDMS/cell-gel system was done using an FEM viscoelastic model developed in COMSOL Multiphysics 6.1. Transient simulations of the PDMS/hydrogel deformation during impact were performed on a 2D-axisymmetric system, assuming symmetry along the impact centerline. Geometry and mesh are illustrated in Figure S5A. The domain thickness is 3 mm (1 mm hydrogel height and 2 mm PDMS height), the radius of the hydrogel is 1.05 mm, and the radius of the PDMS is 4 mm. The PDMS domain radius was substantially larger than the hydrogel to ensure the radial boundaries do not have an effect on the impact location (radius of 0.25 mm) and deformation of the system.

A triangular mesh consisting of 20676 elements was generated for the system, with a minimum element quality of 0.6312 and an average of 0.9459. Mesh refinement around the impactor limited the maximum element size to 0.02 mm, 1/25 the size of the impact radius, and a maximum element growth of 1.005.

A union between the PDMS and hydrogel domains ensures perfect contact during the simulation. The boundary conditions for the simulation include a fixed bottom boundary where the PDMS-hydrogel system sit on the glass petri dish, a free boundary along the right-side and top surface that allow the system to freely deform, an axisymmetric left-side boundary, and a uniform distributed load along the impactor radius to represent the force of the impactor.

The distributed load was applied as a function of time and was derived using a mass-double spring-damper system in eq. 1a and 1b.

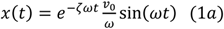

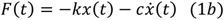

Where x(t) is the displacement function, ζ is the damping ratio, ω = √k/m, v_0_ = √2gh, k is the system stiffness, and c = 2ζmω. k and ζ can be found from material properties for the PDMS and hydrogel, m is the mass of the impactor, h is the height that the mass was dropped, and g is the gravitational constant. Time dependent simulations were run for 50 µs using 1 µs time steps.

*Calcium imaging:* Calcium imaging was performed immediately after injury (0.5hr), 24 hr and 72 hr post-injury. Fluo-4 AM (Thermo Fisher) was dissolved in 20% Pluronic F-127 (in DMSO) and diluted (10 μL/mL) in PFC media to reach a 10 μM final concentration. A loading solution was made by adding 8 μL of stock solution to 1 mL of PFC media. Media in the device was replaced with loading solution and the device was incubated for 60 minutes. The device was transferred back to PFC media for recording. Calcium recordings were acquired using Leica DM-IRBE inverted microscope with a FITC filter (Ex/Em: 488/525 mm). Images from 3 regions of interest (ROIs; B1, B2, and region between the two) were obtained separately for 5 minutes at 20 fps with 50 ms exposure. The device and cells were maintained under physiological conditions [37°C with 5% CO_2_ medical grade air mixture (Airgas, PA)] in a microscope stage top incubator (Tokai hit live cell culture stage adapter) for the entire recording session.

*Neuronal Activity and Network Analysis:* Calcium traces were extracted using custom MATLAB® scripts using a combination of top-hat filter, greedy algorithm, background subtraction and photobleaching correction. For each identified cell body, fluorescence was converted to normalized ΔF/F0 using an estimated baseline F0 (sliding-window percentile) to reduce the impact of slow drift and photobleaching. Deconvolution and spike estimation was performed using MATLAB script developed by Pnevmatikakis ^[19]^. Inferred event trains were used to compute active neurons (AN; defined as ≥5 inferred events/min) and mean firing rate (mFR; events/min). Waveform features (rise/fall time, pulse width, and amplitude) were calculated from ΔF/F0 traces using methods published previously ^[20]^. Because calcium signals integrate over and filter underlying voltage dynamics and depend on dye loading and buffering, spike/event inference provides an indirect estimate of neuronal firing. Therefore, all spike-derived metrics are reported as inferred activity measures for between-group comparisons.

Network analysis was assessed via pair-wise phase-locking synchronization matrix using previously published methods ^[34]^. Briefly, each neuron *j* has a discrete sequence of spike times, *t_j_*(*n*), and a time-varying instantaneous phase was assigned to each neuron j within the nth inter-spike interval (2).

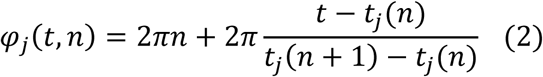

Pair-wise phase-locking synchronization matrix *C* of neurons with total number of M was generated by calculating circular variance of the phase difference between pair-wise neurons *j* and *k* (3).

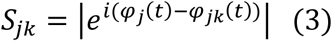

To obtain a normalized value of global synchronization index (GSI), which ranges from 0 (non-coordinated activity) to 1 (perfectly synchronized activity), and independent of the number of neurons, we followed previously published eigen-value based decomposition approaches ^[156, 157]^. The eigenvalue decomposition of C can be given by (4)

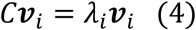

Where *λ_i_* is the eigenvalue and *λ*_1_ ≤ *λ*_2_ ≤ ⋯ ≤ *λ_M_*, and ***v****_i_* is the eigenvector corresponding to *λ_i_*. Surrogate matrices of matrix C were generated using the amplitude-adjusted Fourier transform (AAFT) and repeated 10 times. The mean of eigen-value calculated across 10 surrogates is denoted as ̅*_i_*. The normalized GSI can be computed by (5)

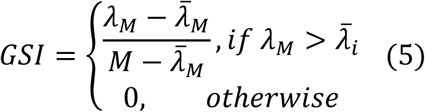

The equation to compute the number of clusters is expressed as (6)

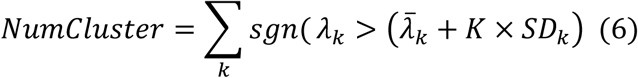

Where *sng* is 1 if 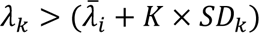 is true, *K* is a constant (here *K* = 2, giving 95% confidence levels), ̅*_k_* is the average and *SD_k_* is the standard deviation of surrogate eigenvalues.

Weighted Modularity (*Q^w^*) describes the degree to which the network may be subdivided into such clearly delineated and nonoverlapping groups, which was measured following previously published method ^[35, 36]^ (7).

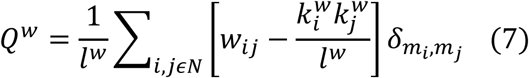

Where *i, j* are the nodes in the network, and N is the set of all nodes in the network. *w_ij_* is the weight of the link (*i,j*); Sum of all weights in the network, *l^w^* = ∑*_i_*_,*jεN*_ *w_ij_*; Weight degree of *i*, *k_i_^w^* = ∑*_jεN_ w_ij_*; *m_i_* is the module containing node *i*, which is determined by Louvain clustering, and *δ_mi_*_,*m*_*_j_* = 1 if *m_i_* = *m_j_*, otherwise *δ_mi_*_,*m*_*_j_* = 0.

Weighted Global Efficiency (*E^w^*) is the average inverse shortest path length. It describes the integration of the network and how efficiently the network exchange information. *E^w^* was calculated following previously published method ^[36, 158]^ (8).

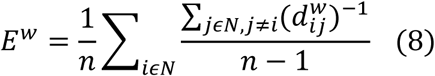

Where *i, j* are the nodes in the network, and *n* is the number of nodes in the network; Shortest weighted path length between *i* and *j*, 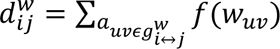, where *f* is a map from weight to length and 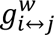 is the shortest weighted path between *i* and *j*.

*Immunocytochemical analysis:* The devices were fixed with 4% (w/v) paraformaldehyde and 0.4 M sucrose in PBS for 20 min at RT. Next, the devices were washed and permeabilized with PBST (0.5% (v/v) TritonX-100 in PBS) and blocked for 1 hr with blocking solution (5% (w/v) BSA in PBST). Primary antibody incubation was performed overnight at 4°C in blocking solution. The devices were then washed with PBST twice and incubated with blocking solution for 1 hr at RT. Secondary antibody incubations were performed for 3 hr at RT in blocking solution, followed by 30 min incubation with Hoechst. Unbound antibodies and Hoechst were washed away by washing 2 times with PBS. Images were acquired on a Zeiss LSM 900 confocal microscope (Zeiss, Germany; Objective: EC Plan-Neofluar 10x/0.30 M27; Detector: GaAsP-PMT). Identical LUTs and acquisition parameters were used across groups within each experiment in the methods section. Detailed antibodies and exposure setting is included in the supplementary method. Images were analyzed using custom MATLAB scripts.

*Western Blotting:* Cells were extracted from hydrogels, washed once with ice-cold DPBS without Ca/Mg, and then lysed in RIPA buffer (ThermoFisher, Waltham) supplemented with 1X Halt Protease & Phosphatase Inhibitor cocktail (Thermo Scientific, Waltham), on ice for 30 min, with intermittent vortexing. Cell lysates were spun down at 14,000 RPM for 20 minutes, at 4⁰C. Supernatants were collected and protein amounts were estimated using Pierce BCA Protein Assay kit (ThermoFisher, Waltham). Sodium dodecyl-sulfate polyacrylamide gel electrophoresis (SDS-PAGE) was run to resolve proteins (10μg per lane) and transferred onto nitrocellulose membranes. The membranes were blocked for 1 hr using 5% non-fat dry milk in TBST and incubated with primary antibodies – Calpain-1 (#MA3-940, Invitrogen), Caspase-3 (#9662, Cell Signaling), pT181 (#MN1050, Thermo), Tau46 (#4019S, CST), pS396 (#ab109390, Abcam), AT8 (#MN1020, Thermo), pT231 (#ab151559, Abcam) and β-actin (#4970S, CST) in 5% BSA in TBST, at 4°C overnight. This was followed by secondary antibody incubation for 1 hr at room temperature. Immunoblots were developed using Clarity Western ECL substrate (170-5060, Bio-Rad) and digital images were acquired using the ChemiDoc MP imaging system (Bio-Rad). Quantification of protein bands was performed using ImageJ software (NIH) and band intensities were normalized to respective β-actin and Tau46 loading controls.

*Secretome Analysis:* Conditioned media was collected before each calcium imaging session at 24 hr, 72 hr, 5d and 8d time points. Pooled replicates were analyzed using 48-plex, human supplemental biomarker 10-plex, and human amyloid beta and tau 4-plex discovery assays (Eve Technologies). Low-abundance proteins were excluded. Proteins with log₂ fold change >0.3 (injury vs. control) at any time point were retained, identifying 24 cytokines and 4 tau-related markers. These log2 fold change values of each marker at each time point were subjected to hierarchical clustering using the Ward algorithm ^[159]^ for their release profile. Next, a cross-correlation matrix of absolute fluorescence values from the 28 markers was used to compute an adjacency matrix after removing non-significant correlations (p-value < 0.5). Functional clusters were visualized using a spectral clustering algorithm, and the functional association of the markers was plotted using a force directed graph. To correlate secretome with neuronal function, sparse canonical correlation analysis (sCCA) was applied to averaged calcium imaging metrics from corresponding biological replicates.

*Statistical Analysis*: The statistical analysis of this study was performed using MATLAB, R Studio and GraphPad Prism 11.0.1 software. Z-score standardization and reference sample normalization were performed to standardize data. Hierarchical clustering and robust regression and outlier removal analysis were performed to remove outliers. Sample size, statistical and post-hoc methods for each statistical analysis were reported in each figure legend. The p-values were reported in the figure. The results were expressed as mean ± standard deviation or mean ± 95% confidence intervals as indicated in each figure legend.

## Supporting information

Legend for Supplementary Movies

## Acknowledgements

This work was supported by National Institute of Health (NIH)-National Institute of Neurological Disorders and Stroke (R01NS099596, and R21NS130468), and Alliance for Regenerative Rehabilitation Research and Training (AR3T) technology development award (to L. K.), and NSF Engineering Research Center (ERC) for Cell Manufacturing Technologies (CMaT, NSF-EEC 1648035) funding (to AGF, SLS, and L. K.). This research was supported in part by NIH R01ES033892 and the Dianne Isakson Distinguished Professorship award (to J. R. R.).

## Data Availability Statement

A full set of raw data is attached to the supporting information. Computer-aided design (CAD) files for the Neuron-on-Chip device, and raw images (calcium imaging, immunocytochemistry) will be made available by the lead contact upon reasonable request. Associated MATLAB code for calcium imaging data processing is available through https://github.com/Thorowen/Calcium-imaging-data-processing.

## Author contributions

R. T., C-F L., and L. K. designed the study. R. T., C. K., and L. K. wrote the manuscript. R. T., M. M. S., and H. W. performed neuronal differentiation and cell culture. R. T., C-F L. performed device design and manufacturing. R. T., and A. C. performed, injury induction, calcium imaging, and immunofluorescence staining experiments. M. M. S., N. G., and C. K. performed western blotting. A. L. performed stiffness measurement and dynamic mechanical analysis. J. S. performed finite element modeling analysis. R. T. and A. M. performed data analysis. J. S., I. M-W., A. F., J. P., N. Z, J. R, C-F L, S. L. S. and L.K. assisted with overall experimental design and manuscript revisions.

## Declaration of interests

The authors declare no conflicts of interest.

Received: ((will be filled in by the editorial staff))

Revised: ((will be filled in by the editorial staff))

Published online: ((will be filled in by the editorial staff))

