## Supplementary Figures for "A 3D Human Neuron-on-Chip Platform to Monitor Neuronal Injury Responses"

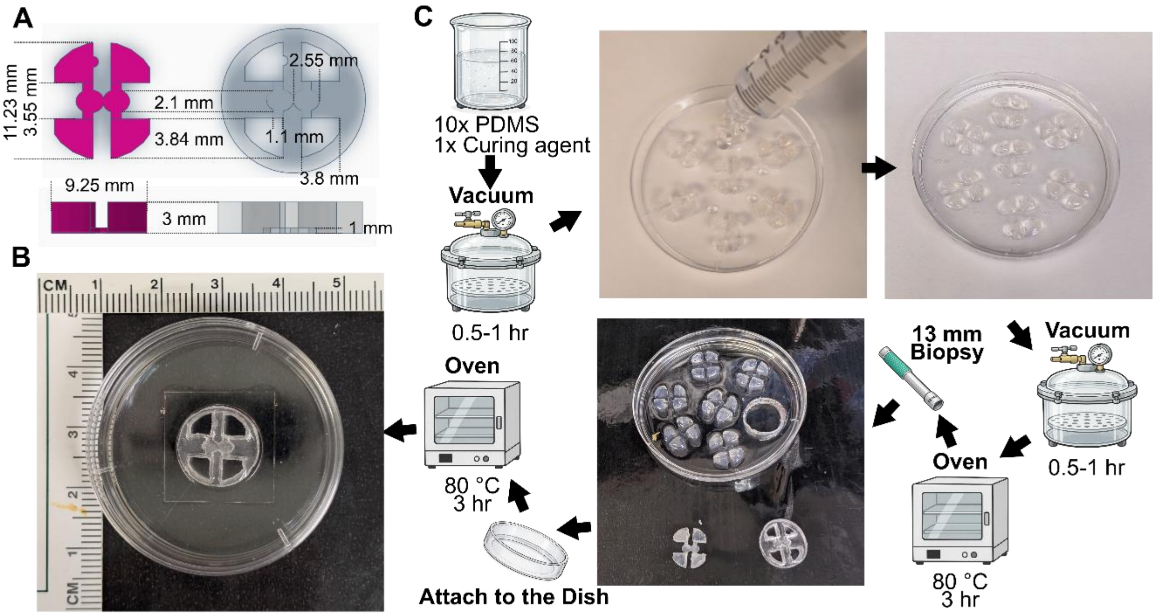


**Figure S1.** Neuron-on-Chip device dimensions. (A) Top and front views of 3D printed Neuron-on-Chip 3D mold and PDMS shells with dimension measures. (B) Photograph of fully assembled Neuron-on-Chip device. (C) Fabrication procedure of Neuron-on-Chip device.

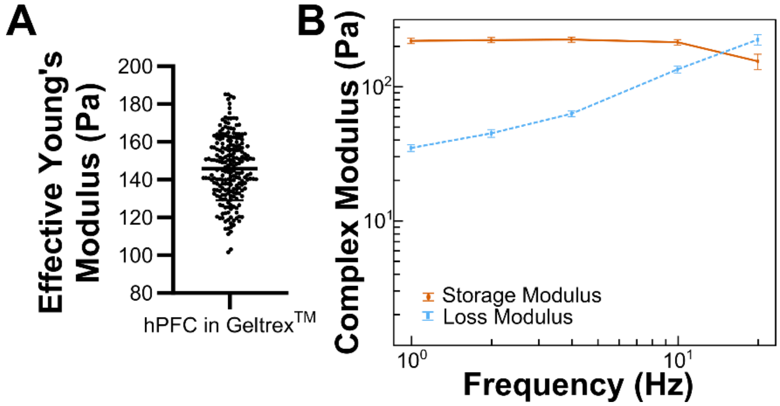


**Figure S2.** Characterization of the physical properties of cell-gel construct. (A) Measurement of effective Young's modulus of hPFC/Geltrex™ complex. Error bars indicate mean ± standard deviation. (B) Results from the dynamic mechanical analysis reveal the crossover of the elastic and viscoelastic behavior of the cell-gel construct at around 10-20 Hz oscillation. Error bars indicate mean ± standard deviation.

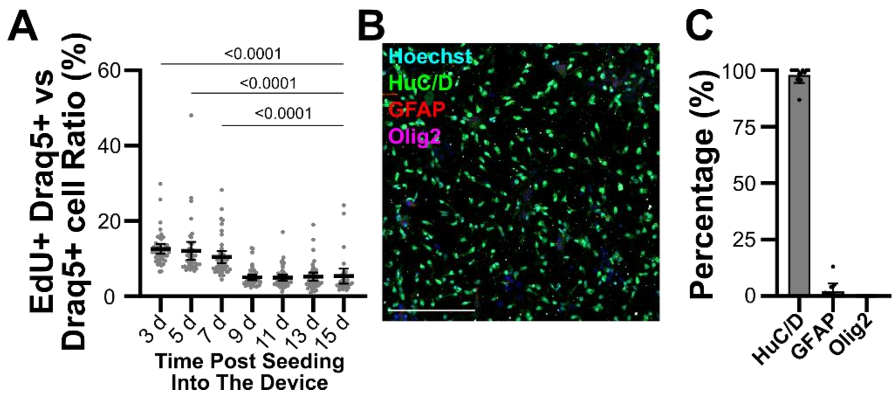


**Figure S3.** hPFC cells demonstrate decreased proliferation and increased neuronal differentiation post device-seeding. (A) EdU+ staining of hPFC cells reveals significantly reduced cell proliferation 9-days post device seeding. Error bars indicate mean ± standard deviation. One-way ANOVA with Tukey post-hoc. Significant p-value (<0.05) of pair-wise comparisons with 15 d are reported in the figure. n = 5 images per sample, data represents 3 independent experiments. (B) Representative confocal image of immunohistochemically stained HuC/D (pan-neuronal marker, green), GFAP (astrocyte marker, red), Olig2 (oligodendrocytes marker, magenta), and Hoechst (blue), scale – 200 µm. (C) Quantification of (B) demonstrating a major presence of HuC/D+ neurons in PFC culture 2-weeks post device seeding. Error bars indicate mean ± standard deviation. n = 5 images per sample from 3 independent experiments.

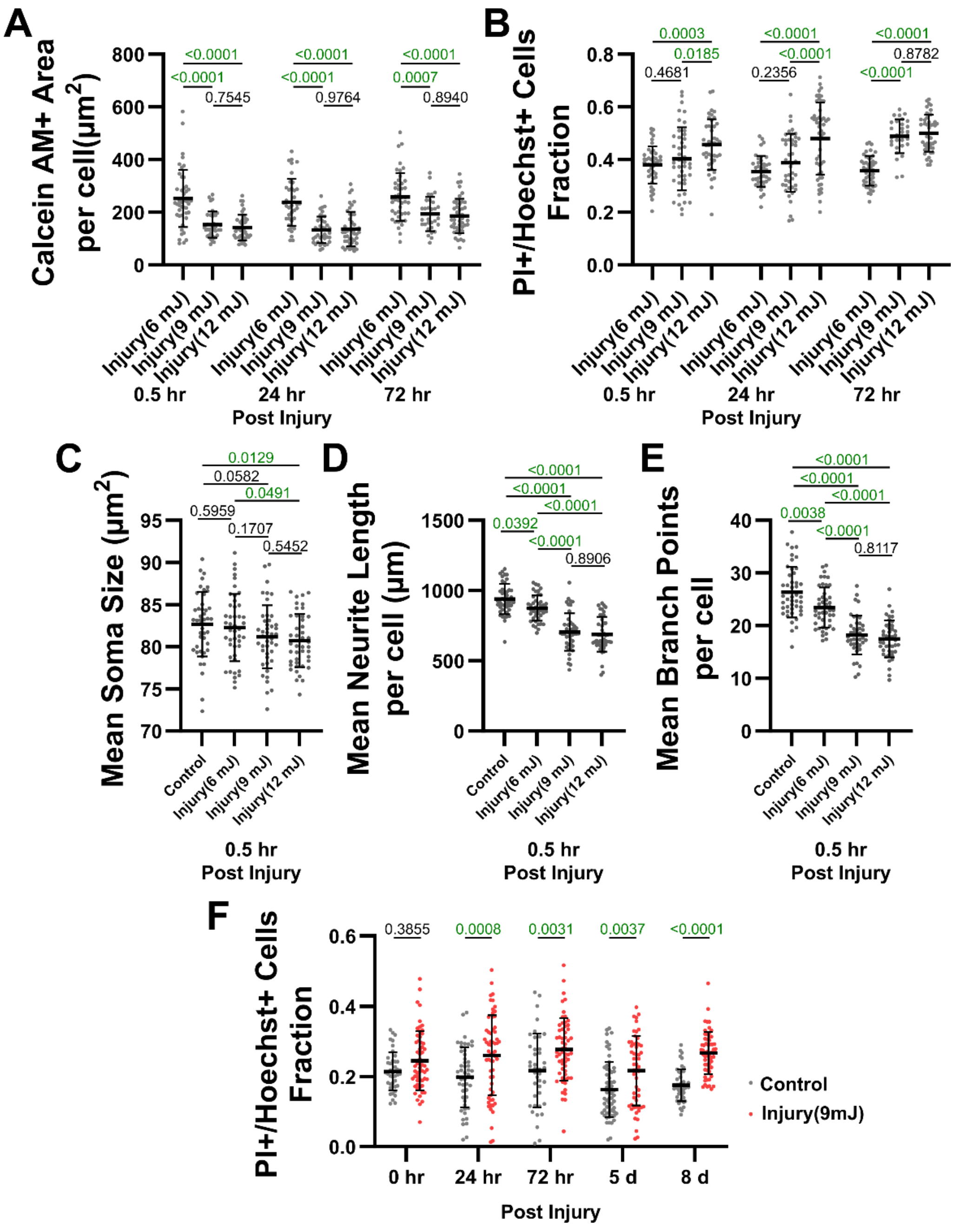

**Figure S4.** Weight-drop injury induces changes in cell morphology and morbidity in response to increasing impact force. (A-B) Scatter plots demonstrating significant changes in cell body retraction (A), and cell death (B) with increasing impact force. Error bars indicate mean ± standard deviation. Two-way ANOVA with Tukey post-hoc. P-value of pair-wise comparisons between groups at each tested timepoint are reported in the figure and significant p-value (<0.05) is highlighted in green. (C-E) Scatter plots demonstrating significant changes in neuronal soma size (C), neurite length (D), and branch points number (E) with increasing impact force. Error bars indicate mean ± standard deviation. One-way ANOVA with Tukey post-hoc. P-value of pair-wise comparisons are reported in the figure and significant p-value (<0.05) is highlighted in green. (F) Scatter dot plot demonstrating significantly enhanced cell death in the weight drop injured group, compared to uninjured control. Error bars indicate mean ± standard deviation. Two-way ANOVA with Bonferroni post-hoc. . P-value of pair-wise comparisons between injury vs control groups at each tested timepoint are reported in the figure and significant p-value (<0.05) is highlighted in green. 10 images per device, 3 devices per group from 3 independent experiments.

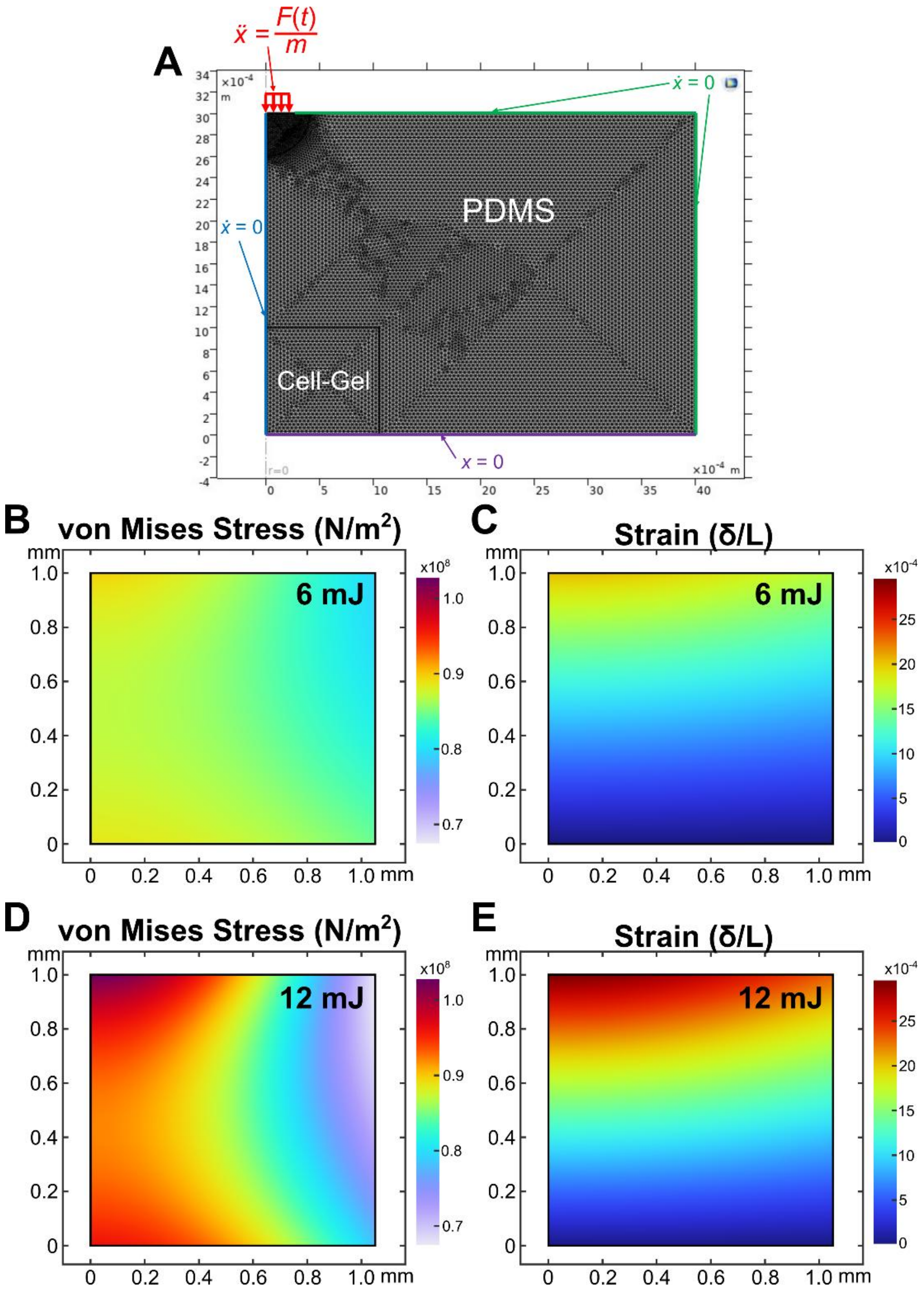


**Figure S5.** Finite element modeling (FEM) revealing changes in stress/strain to cell-gel in response to increasing impact energy. (A) 2D-axisymmetric geometry and mesh of the PDMS-hydrogel system. The left boundary illustrates the symmetrical condition (blue), the bottom boundary is fixed (purple) and the right and top boundaries can freely displace (green). A uniform distributed load (red) represents the impactor being dropped onto the test region for the transient viscoelastic FEM model. (B) Heatmaps of the von Mises Stress and surface strain of the cell-gel construct under 6 mJ (B, C) amd 9 mJ (D, E) injury condition.

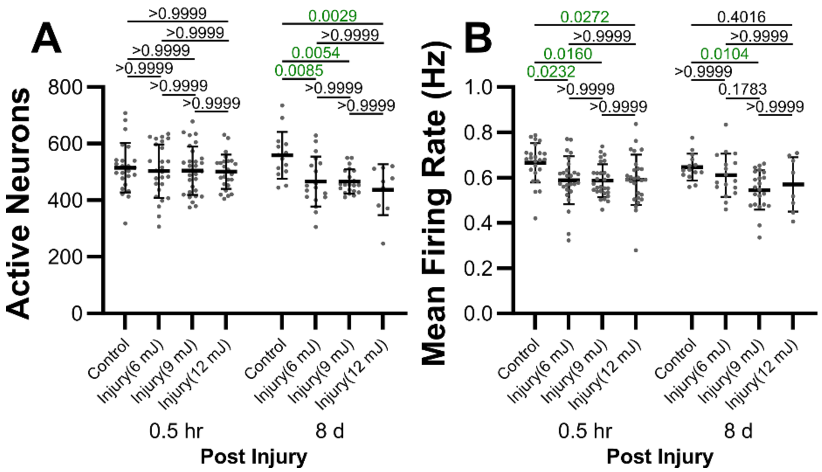


**Figure S6.** Neuronal activity level across injury conditions. (A, B) Scatter plots reveal no significant changes in active neuron number (A), and mean firing rate (B) across two injury conditions 0.5 hr and 8 d post injury. Plot shown in mean±sd. Two-way ANOVA with Bonferroni post-hoc. P-value of pair-wise comparisons between groups at each tested timepoint are reported in the figure and significant p-value (<0.05) is highlighted in green. 3 ROIs per device, 3 devices per time point from 3 independent experiments.

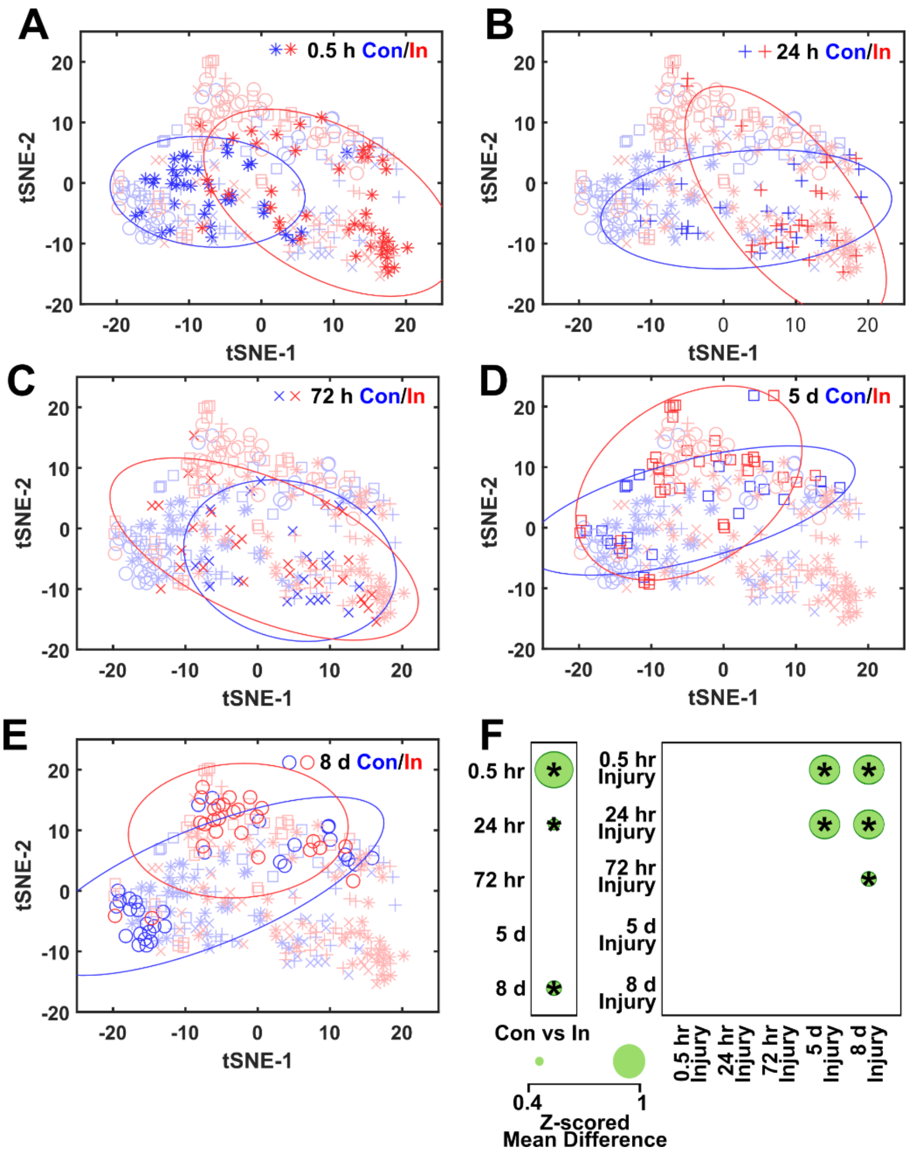


**Figure S7.** Injury induces significant changes in neuronal activity at 0.5h, 24h, and 8d post injury. (A-E) tSNE plot of sample distribution and their 95% CI ellipses on neuronal functions described by 13 neuronal activity parameters. (F) Bubble plots reveal significant differences in neuronal activity pattern between injury and control groups at 0.5 hr, 24 hr, and 8 d post injury, and between early (0.5-24 hr) and late (5-8 d) injury responses. PERMANOVA, P-Group = 0.001, iterations = 1000. Bonferroni pairwise comparison, *p = 0.0450. Blurred markers in each plot indicate samples from other time points.

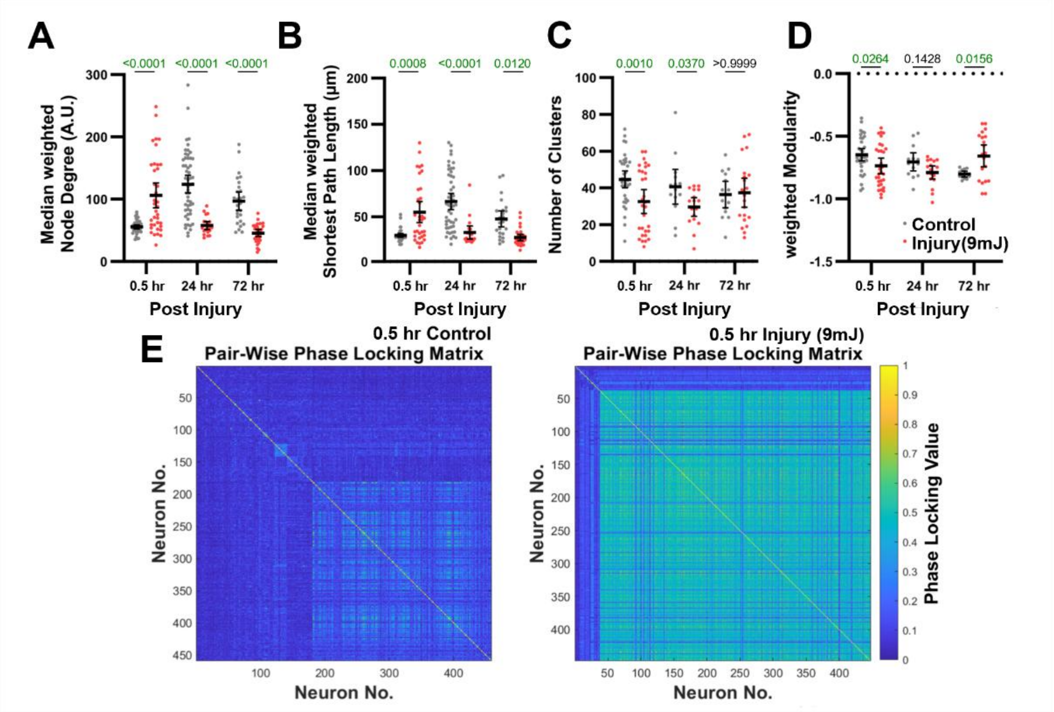


**Figure S8.** Injury induces an acute increase in neuronal connectivity and disruption of network structure. (A, B) Median weighted Node Degree and median weighted Shortest Path Length indicate acute increase and subsequent loss of functional connections. (C, D) NoC (C, neuronal subcommunities) identified by an eigen-value-based algorithm, and Modularity (D, sub-community presence) demonstrate acute disruption of community structure in response to injury. (E) Representative heatmaps of pair-wise phase locking matrix from control and injured groups 0.5 hr post injury indicate remodeling of functional network structure in response to the injury. Error bars indicate mean ± 95% confidence interval. Two-way ANOVA with Bonferroni post hoc. P-value of pair-wise comparisons between injury vs control groups at each tested timepoint are reported in the figure and significant p-value (<0.05) is highlighted in green. 3 ROIs per device, 3-5 devices per time point from 4 independent experiments.

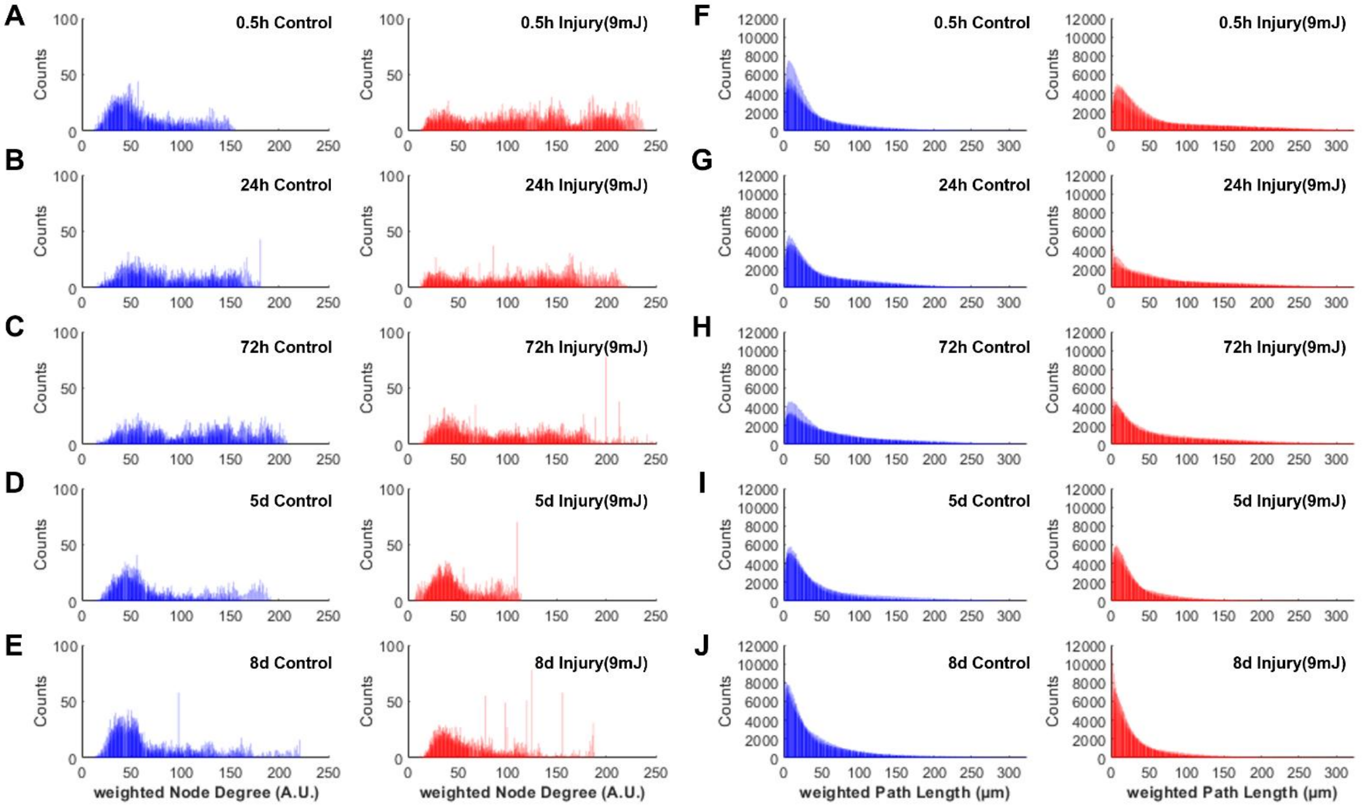


**Figure S9.** Loss of overall connectivity and long-range connections post injury. Histograms demonstrate the loss of neurons with high connectivity (A-E) and long-range connection (F-J) over time post-injury in the injured groups when compared to controls.

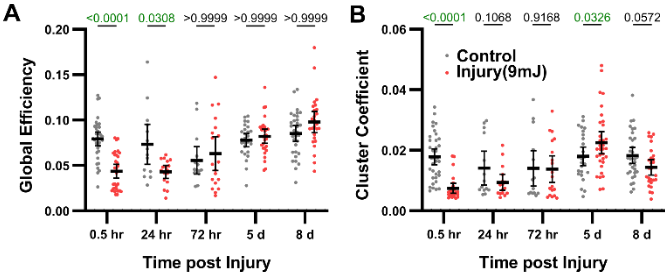


**Figure S10.** Acute decrease in global efficiency and cluster coefficient post-injury. (A-B) Network analysis demonstrated an injury induced acute alteration of Global Efficiency (A), and Cluster Coefficient (B). Error bars indicate mean ± 95% confidence interval. Two-way ANOVA with Bonferroni post hoc. P-value of pair-wise comparisons between injury vs control groups at each tested timepoint are reported in the figure and significant p-value (<0.05) is highlighted in green. 3 ROIs per device, 3-5 devices per time point from 4 independent experiments.

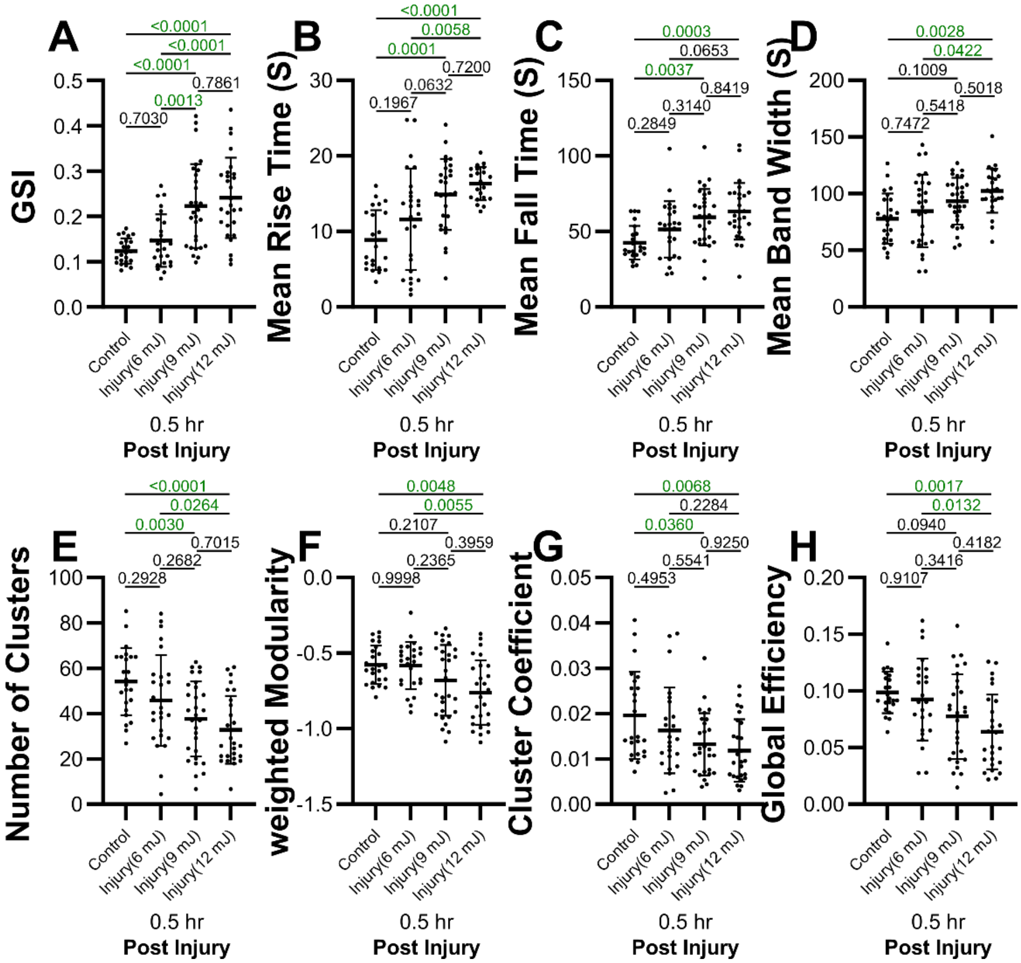

**Figure S11.** Stress-dependent early post injury response. (A-H) Multiple signal-cell calcium dynamic, network structure and function parameters reveal stress-dependent injury responses at 0.5 hr post injury. Error bars indicate mean ± standard deviation. One-way ANOVA with Tukey post hoc. P-value of pair-wise comparisons between injury vs control groups at each tested timepoint are reported in the figure and significant p-value (<0.05) is highlighted in green. 3 ROIs per device, 3 devices per time point from 3 independent experiments.

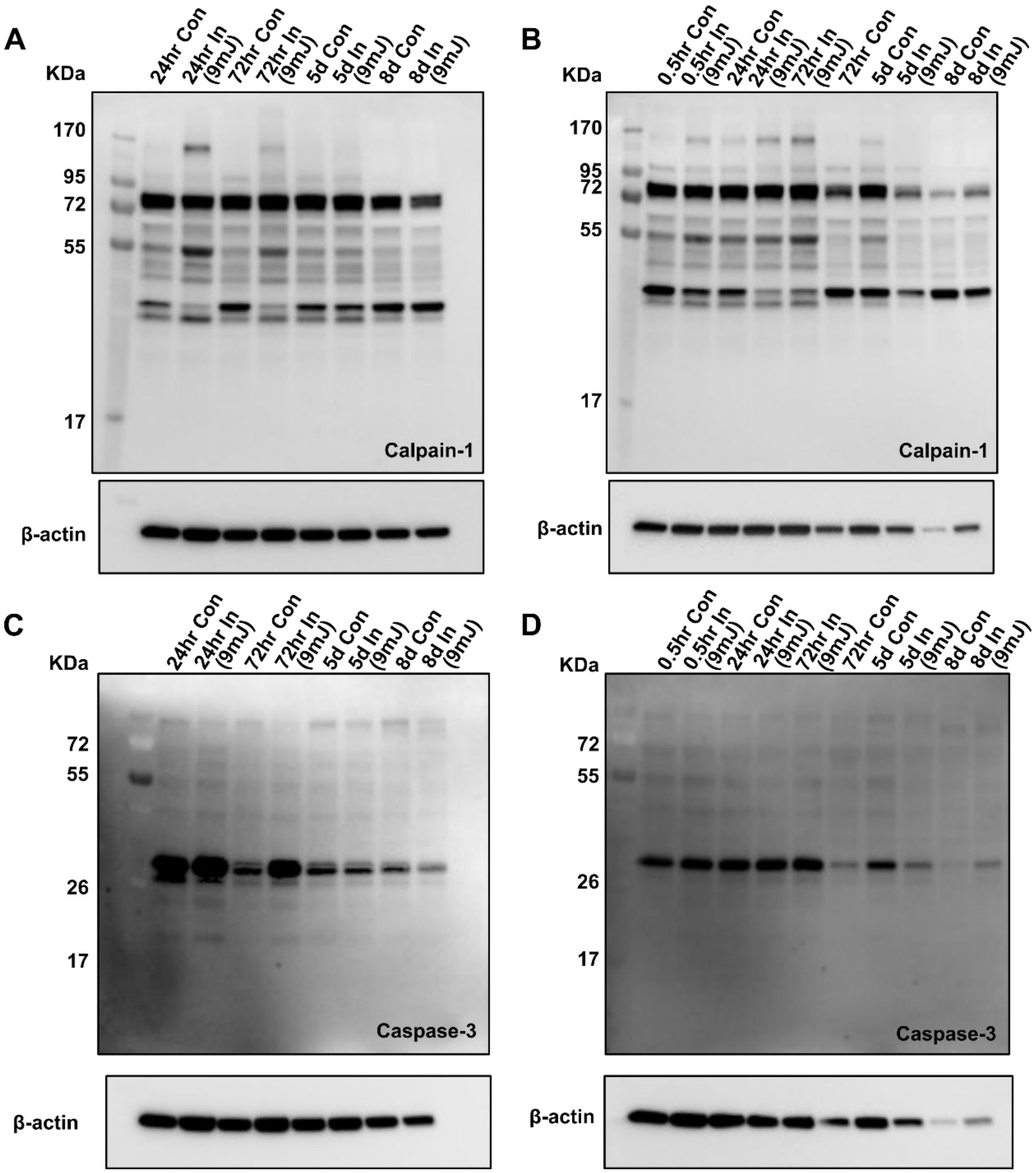


**Figure S12.** Western blot analysis of calpain-1 and caspase-3. (A, B) Western blot showing calpain-1 bands at 0.5 hr – 8 d post injury. (C, D) Western blots showing caspase-3 bands at 0.5 hr – 8 d post injury.

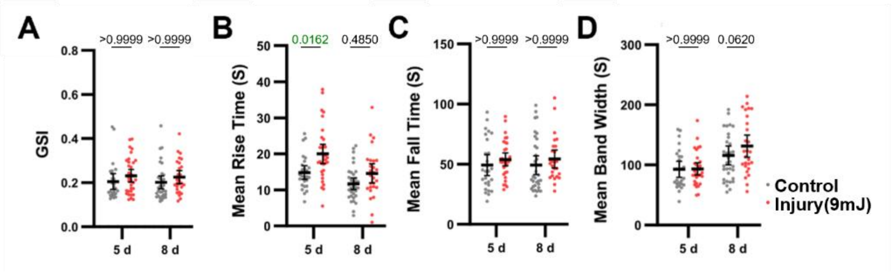


**Figure S13.** Dysregulation of calcium trafficking 5-8 d post injury. GSI (A), mean Rise Time (B), mean Fall Time (C), mean Band Width (D) revealing a statistically insignificant but increasing trend in injured neurons 5-8 d post injury compared to control. Error bars indicate mean ± 95% confidence interval. Two-way ANOVA with Bonferroni post hoc. P-value of pair-wise comparisons between injury vs control groups at each tested timepoint are reported in the figure and significant p-value (<0.05) is highlighted in green. 3 ROIs per device, 3-5 devices per time point from 4 independent experiments.

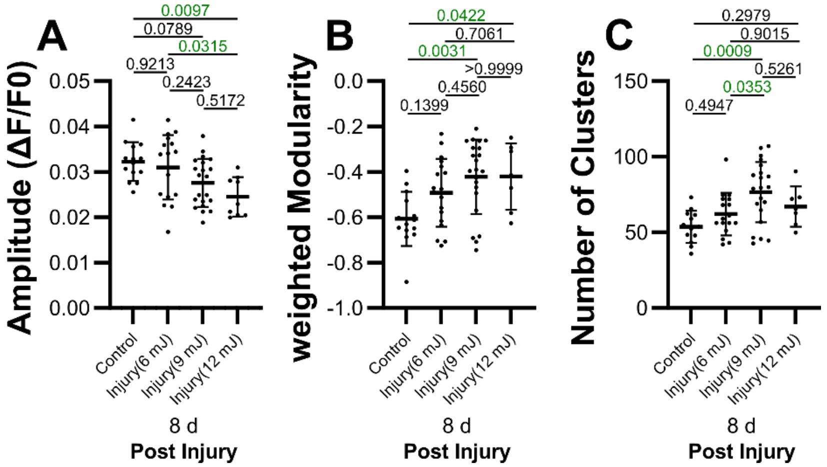


**Figure S14.** Late injury response across injury conditions. (A) Injury induced reduction in Amplitude that scales proportionately with impact energy. (B, C) Enhanced network fragmentation indicated by community structure parameters - Modularity (B) and Number of Clusters (C). Error bars indicate mean ± standard deviation. One-way ANOVA with Tukey post hoc. P-value of pair-wise comparisons between injury vs control groups at each tested timepoint are reported in the figure and significant p-value (<0.05) is highlighted in green. 3 ROIs per device, 2-3 devices per time point from 3 independent experiments.

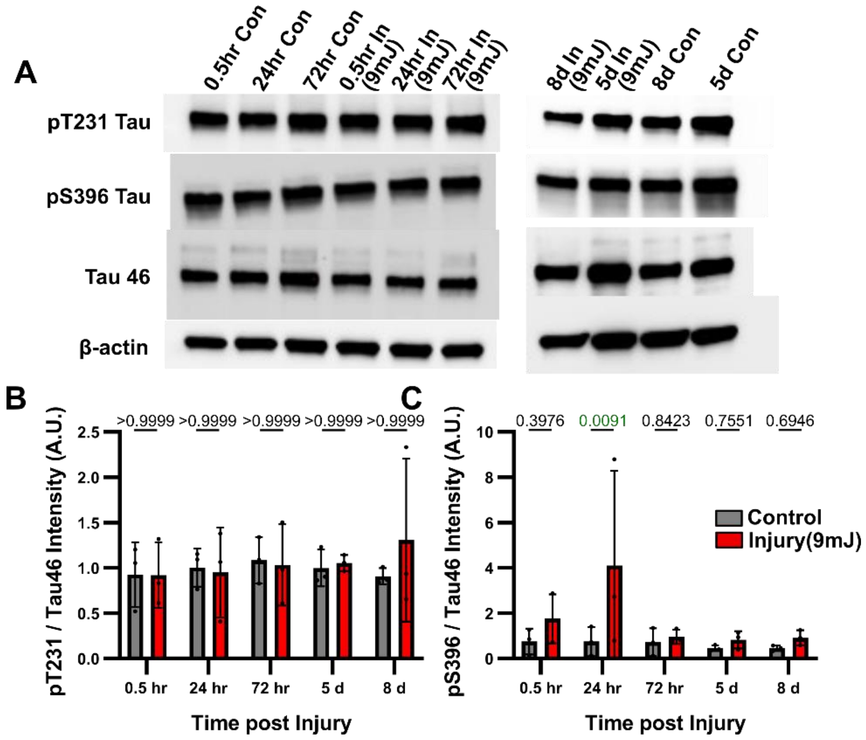


**Figure S15.** Intracellular accumulation of pathologically relevant tau. (A) Western blotting results of pT231, pT396 tau and total tau levels. (B-C) Densitometric quantification of results from A. Error bars indicate mean ± standard deviation. Two-way ANOVA with Bonferroni post hoc. P-value of pair-wise comparisons between injury vs control groups at each tested timepoint are reported in the figure and significant p-value (<0.05) is highlighted in green. 3-4 devices per sample, 1 sample per time point from 3 independent experiments.

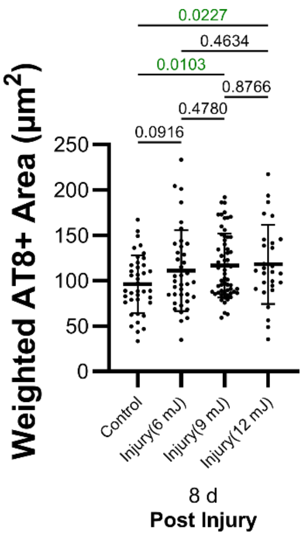


**Figure S16.** AT8+ Tau upregulation post injury across injury conditions. Error bars indicate mean ± standard deviation. One-way ANOVA with Tukey post hoc. P-value of pair-wise comparisons between injury vs control groups at each tested timepoint are reported in the figure and significant p-value (<0.05) is highlighted in green. 8-10 images per device, 2-3 devices per group, from 2 independent experiments.
