## Supplementary Method for "A 3D Human Neuron-on-Chip Platform to Monitor Neuronal Injury Responses"

### Supplementary Methods

*Cell Culture of hPSC-Derived Prefrontal Cortex Neurons:* Human pluripotent stem cells (hPSCs) derived prefrontal cortex neurons (hPFCs) were generated following methods described previously [1] using Mel1 cell line (NIHhESC-11-0139, passage number 40-50). hPSCs were dissociated into single cells and plated on Matrigel-coated plates in Essential 8 medium with 10 µM ROCK inhibitor. From day 0–2, cultures were treated with Essential 6 medium containing 2 µM XAV939, 100 nM LDN193189, and 10 µM SB431542. XAV939 was withdrawn from day 2 onward. At day 6–8, cells were replated as high-density droplets on poly-ornithine/laminin/fibronectin-coated dishes and cultured in N2 medium with B27 (1:1000, no vitamin A), FGF8 (50 ng/mL), and SHH (25 ng/mL) for 4 days, until neuroepithelial rosettes were visible. Droplets were then passaged 1:2 with trypsin onto poly-ornithine/laminin/fibronectin coated plates and cultured in the same media. On day 16, cells were passaged using Accutase and plated at $1\times10^6$ cells/cm$^2$ in PFC medium (Neurobasal™, B27 1:50, N2 1:100, GlutaMAX 1:100). FGF8 (50 ng/mL) was supplemented during the first two weeks. Cells were passaged again on day 22, replated at 100,000 cells/cm$^2$ and used for experiments on day 35.

*Immunocytochemical analysis:* Immunofluorescent images were acquired on a Zeiss LSM 900 confocal microscope (Zeiss, Germany; Objective: EC Plan-Neofluar 10x/0.30 M27; Detector: GaAsP-PMT). Antibodies used for the studies are listed as follows:

Primary antibodies include: pS396 Tau (Abcam #ab109390), pT231 Tau (Abcam #ab151559), Tau 46 (Cell Signaling #4019S), AT8 (ThermoFisher #MN1020), Beta III-

Tubulin (EMD Millipore #ab9354), Synaptophysin-1 (Abcam #ab14692), MOAB2 (Abcam #ab126649). Secondary antibodies include: 488 Goat anti-Chicken IgY (ThermoFisher #A11039), 594 Goat anti-Rabbit IgG (H+L) (ThermoFisher #A21428), 647 Goat anti-Mouse IgG1 (ThermoFisher #20240), 647 Goat anti-Rabbit IgG (H+L) (ThermoFisher #21244).

Exposure settings used for each confocal panel are listed below:

Hoechst/B3T/F-Actin/Syn-1 panel: Hoechst – 405 nm: 1.50%, Detector Gain: 778 V; B3T – 488 nm: 3.10%, Detector Gain: 817 V; F-Actin – 561 nm: 3.60%, Detector Gain: 836 V; Syn-1 – 640 nm: 3.0%, Detector Gain: 749 V. Hoechst/HuCD/GFAP/Olig2 panel: Hoechst – 405 nm: 0.80%, Detector Gain: 856 V; HuCD – 488 nm: 0.60%, Detector Gain: 738 V; GFAP – 561 nm: 0.80%, Detector Gain: 772 V; Olig2 – 640 nm: 1.0%, Detector Gain: 704 V. Hoechst/AT8 panel: Hoechst – 405 nm: 1.60%, Detector Gain: 863 V; AT8 –nm: 3.60%, Detector Gain: 768 V. Hoechst/Fluo-4 AM/NFT panel: Hoechst – 405 nm: 2.80%, Detector Gain: 864 V; Fluo-4 AM – 488 nm: 2.70%, Detector Gain: 772 V; NFT – 561 nm: 3.50%, Detector Gain: 720 V. Hoechst/Fluo-4 AM/MOAB2 panel: Hoechst – 405 nm: 2.80%, Detector Gain: 864 V; Fluo-4 AM – 488 nm: 2.70%, Detector Gain: 772 V; MOAB2 – 640 nm: 0.50%, Detector Gain: 782 V.
