## Supplementary material for "A 3D Human Neuron-on-Chip Platform to Monitor Neuronal Injury Responses": Legend for Supplementary Movies

### Legends for supplementary movies

**Movie S1**. Visualization of neuronal morphology in the Neuron-on-Chip device before injury with 3D reconstruction. (blue: Hoechst – nuclei, green: Calcein AM – cell body).

**Movie S2**. Visualization of neuronal morphology in the Neuron-on-Chip device 0.5 hr after injury (9 mJ) with 3D reconstruction. (blue: Hoechst – nuclei, green: Calcein AM – cell body).

**Movie S3**. Representative calcium imaging recording from control group 0.5 hr post injury (9 mJ). Accelerated 30 times. Scale bar - 100 μm. (green: Fluo4-AM).

**Movie S4**. Representative calcium imaging recording from injury group 0.5 hr post injury (9 mJ). Accelerated 30 times. Scale bar - 100 μm. (green: Fluo4-AM).

**Movie S5**. Representative calcium imaging recording from control group 8 d post injury (9 mJ). Accelerated 30 times. Scale bar - 100 μm. (green: Fluo4-AM).

**Movie S6**. Representative calcium imaging recording from injury group 8 d post injury (9 mJ). Accelerated 30 times. Scale bar - 100 μm. (green: Fluo4-AM).
